# Loss of *N*^1^-Methylation of G37 in tRNA Induces Ribosome Stalling and Reprograms Gene Expression

**DOI:** 10.1101/2021.05.27.446073

**Authors:** Isao Masuda, Jae-Yeon Hwang, Thomas Christian, Sunita Maharjan, Fuad Mohammad, Howard Gamper, Allen R. Buskirk, Ya-Ming Hou

## Abstract

*N*^1^-methylation of G37 is required for a subset of tRNAs to maintain the translational reading-frame. While loss of m^1^G37 increases ribosomal +1 frameshifting, whether it incurs additional translational defects is unknown. Here we address this question by applying ribosome profiling to gain a genome-wide view of the effects of m^1^G37 deficiency on protein synthesis. Using *E. coli* as a model, we show that m^1^G37 deficiency induces ribosome stalling at codons that are normally translated by m^1^G37-containing tRNAs. Stalling occurs during decoding of affected codons at the ribosomal A site, indicating a distinct mechanism than that of +1 frameshifting, which occurs after the A site. Enzyme and cell-based assays show that m^1^G37 deficiency reduces tRNA aminoacylation and in some cases peptide-bond formation. We observe changes of gene expression in m^1^G37 deficiency similar to those in the starvation-induced stringent response, consistent with the notion that loss of tRNA aminoacylation activates the response. This work demonstrates a previously unrecognized function of m^1^G37 that emphasizes its role throughout the entire elongation cycle of protein synthesis, providing new insight into its essentiality for bacterial growth and survival.

## Introduction

*N*^1^-methylation of G37 in tRNA, generating m^1^G37 on the 3’-side of the anticodon, is a post-transcriptional modification that is conserved in all domains of life (Bjork *et al*, 2001). It has been specifically associated across evolution with all isoacceptors of tRNA^Pro^, which are species that share the same prolyl specificity of aminoacylation but differ in the primary sequence and in the anticodon triplet and yet collectively decode the Pro CCN codons, where N = A, C, G, and U. Similarly, m^1^G37 has also been conserved in the CCG isoacceptor of tRNA^Arg^ (tRNA^Arg^(CCG), anticodon CCG for pairing with the CGG codon), and in the tRNA^Leu^(GAG), tRNA^Leu^(UAG), and tRNA^Leu^(CAG) isoacceptors (for pairing with CUU and CUC codons (CU[C/U]) and CU[A/G] codons, respectively) (Bjork *et al*., 2001; Li *et al*, 1997), although it may be also present in additional tRNAs in higher eukaryotes. The m^1^G37 methylation of tRNA is essential for cell growth and viability; elimination of the enzyme responsible for the methylation causes cell death in yeast and in bacteria (Bjork *et al*, 2007; Bjork *et al*., 2001; Masuda *et al*, 2019). The established function of m^1^G37 is to maintain the translational reading-frame during protein synthesis (Bjork *et al*, 1989; Hagervall *et al*, 1993; Qian *et al*, 1998). Loss of m^1^G37 elevates frequencies of ribosomal +1 frameshifting in kinetic assays with reconstituted *E. coli* ribosomes *(Gamper et al, 2015a)*, and in cell-based assays in *E. coli* and *Salmonella (Gamper et al, 2021; Gamper et al., 2015a)*. Unlike ribosomal miscoding, +1 frameshifting is almost always deleterious, altering the translational reading-frame, inducing pre-mature termination of protein synthesis, and ultimately cell death.

Recent work has shed light on how loss of m^1^G37 induces ribosomal +1 frameshifting. While non-methylated tRNAs were thought to pair incorrectly in the ribosomal A site (the aminoacyl-tRNA (aa-tRNA) binding site) during decoding (Roth, 1981), they were found to recognize cognate codons in the correct reading-frame in X-ray crystal structures and in kinetic assays (Gamper *et al*., 2021; Maehigashi *et al*, 2014). Maintenance of the correct reading-frame in the A site is consistent with the strict ribosomal A-site structure that selects for accurate anticodon-codon pairing during decoding (Ogle *et al*, 2001; Ogle & Ramakrishnan, 2005). Even genetically isolated high-efficiency +1-frameshifting tRNAs, which usually contain an extra nucleotide inserted to the anticodon loop (Atkins & Bjork, 2009), were found to occupy the A site in the correct triplet reading-frame (Dunham *et al*, 2007; Fagan *et al*, 2014; Gamper *et al*., 2021; Maehigashi *et al*., 2014). These genetically isolated +1-frameshifting tRNAs, however, were found to occupy the triplet +1-frame in the ribosomal P site (the peptidyl-tRNA binding site) (Hong *et al*, 2018).

All existing evidence points to tRNA +1 frameshifting occurring after decoding at the A site. A kinetic study of a frameshift-prone tRNA, harboring the native anticodon loop but lacking m^1^G37, indicates that it can undergo +1 frameshifting in one of the two subsequent steps from the A site – during translocation from the A site to the P site, or during occupancy in the P site next to an empty A site (Gamper *et al*., 2015a). The possibility of +1 frameshifting during translocation is supported by a more elaborate kinetic study of a tRNA with an expanded anticodon loop (Gamper *et al*., 2021). Similarly, the possibility of +1 frameshifting within the P site is supported by X-ray crystal structures of a P-site bound frameshift-prone tRNA with a natural anticodon loop (Hoffer *et al*, 2020). In these structures, while m^1^G37 stabilizes the tRNA on an mRNA codon that is prone to frameshifting, loss of m^1^G37 destabilizes the tRNA-ribosome interaction and induces +1 frameshifting within large conformational changes of the ribosome.

Here we seek to determine whether m^1^G37 plays additional roles beyond maintaining the ribosome translational reading-frame. An earlier study suggested that loss of m^1^G37 might have delayed the tRNA anticodon-codon pairing interaction during decoding (Li *et al*., 1997), raising the possibility of a role at the ribosomal A site. To test this possibility more broadly, we employ the approach of ribosome profiling to determine ribosome positions during translation of the entire transcriptome of a cell (Ingolia *et al*, 2009) and to monitor how ribosome density changes upon loss of m^1^G37. Using *E. coli* as a model, we show that deficiency of m^1^G37 induces global ribosome stalling, most notably at Pro codons CCN, the Arg codon CGG, and the Leu codon CUA, all of which are translated by tRNAs that are normally methylated with m^1^G37. Stalling is most prominent when the affected codons are being decoded at the ribosomal A site, indicating a distinct mechanism than that of +1 frameshifting. Enzyme- and cell-based assays show that m^1^G37 deficiency reduces aminoacylation of the affected tRNAs, including all isoacceptors of tRNA^Pro^ and the isoacceptor tRNA^Arg^(CCG), and that additionally it reduces peptide-bond formation for some of these tRNAs. Most significantly, stalling induces programmatic changes in gene expression that are consistent with changes occurring during the bacterial stringent response under environmental stress of nutrient starvation. These findings support a model in which m^1^G37 deficiency reduces levels of aa-tRNAs at the ribosomal A site and prevents peptide-bond formation in some cases, leading to ribosome stalling. Binding of uncharged tRNAs to the A site would then induce programmatic changes of gene expression similar to those occurring during the bacterial stringent response through activation of the ppGpp synthase RelA (Gourse *et al*, 2018). The importance of m^1^G37 for the ribosomal activity at the A site, together with its already demonstrated importance in maintaining the translational reading-frame from the A site to the P site and within the P site (Gamper *et al*., 2021; Gamper *et al*., 2015a), establishes the involvement of the methylation throughout the entire elongation cycle of protein synthesis. This sustained involvement of m^1^G37 during protein synthesis underscores its indispensable role in bacterial viability and survival.

## Results

### *E. coli* strains with conditional m^1^G37 deficiency

Because m^1^G37 is essential for cell viability, a simple knock-out of the gene responsible for its biosynthesis cannot be made. Previous studies of cellular functions of m^1^G37 relied on temperature-sensitive variants of the gene responsible for m^1^G37 biosynthesis whose protein product became inactivated at elevated temperatures (Bjork & Nilsson, 2003; Masuda *et al*, 2013). Because elevated temperatures induce changes in gene expression, we took a different approach to conditionally deplete m^1^G37 to study its role in protein synthesis. Interestingly, while m^1^G37 is conserved in evolution, the genes responsible for its biosynthesis are distinct – being *trmD* in bacteria and *trm5* in archaea and eukaryotes (Christian *et al*, 2004), and the protein product of each is fundamentally different in structure and mechanism (Christian & Hou, 2007; Christian *et al*, 2010a; Christian *et al*, 2010b; Christian *et al*, 2016; Lahoud *et al*, 2011; Sakaguchi *et al*, 2014). We recently constructed conditional m^1^G37-deficient strains of *E. coli* and *Salmonella*, in which the *trmD* locus is deleted from the chromosome and cell viability of the *trmD-knock-out* (*trmD-KO*) strain is maintained by a plasmid-borne human *trm5* that is under arabinose (Ara)-controlled expression (Gamper *et al*., 2015a; Masuda *et al*., 2019). Upon induction with Ara, expression of human *trm5* is sufficient to supply m^1^G37-tRNAs to support bacterial viability (*trmD-KO (trm5*+)) (Christian *et al*., 2004), whereas upon replacement of Ara with glucose (Glc), expression of human *trm5* is arrested and the human enzyme is degraded inside bacterial cells (*trmD-KO (trm5–*)) (Christian *et al*, 2013). As a control, a *trmD-wild-type* (*trmD-WT*) strain was created, where expression of the plasmid-borne *trm5* in the presence of Ara (*trmD-WT* (*trm5*+)), or its repression in the presence of Glc (*trmD-WT* (*trm5–*)), did not affect cell viability.

To avoid the possibility of artifacts by studying only one conditional m^1^G37-deficient strain, we developed a second conditional m^1^G37-deficient strain to compare data and to strengthen conclusions. In this second conditional strain, we chose *E. coli* as a model and extended *trmD* at the chromosomal locus with a degron sequence, adding YALAA to the C-terminus of the TrmD enzyme (the *trmD-deg* strain) and allowing the protease ClpXP to target the enzyme for rapid degradation. We controlled TrmD degradation by inducing the expression of *clpXP* from a plasmid using Ara, or by repressing the expression using Glc (Carr *et al*, 2012). Because over-expression of the plasmid-borne *clpXP* could also target proteins without the degron tag, we generated a control strain (*trmD-cont*) for comparison to remove non-specifically affected proteins. In this *trmD-cont* strain, we added the coding sequence for the degron tag after the stop codon of *trmD* to maintain the same gene length as in *trmD-deg*, but without expression of the tag (*Figure 1A and Figure 1–figure supplement 1A*).

**Figure 1.**
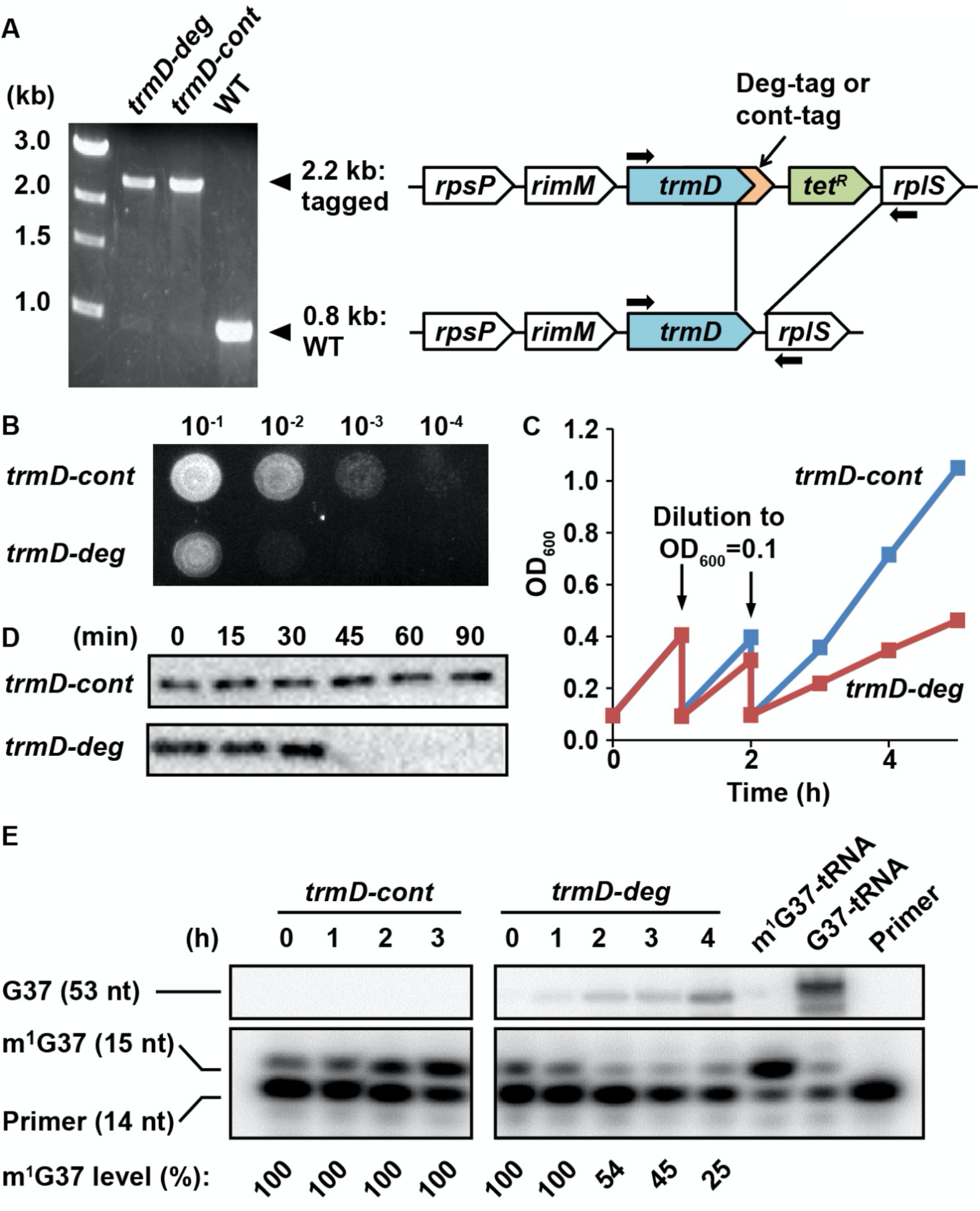
A degron strategy to create conditional m^1^G37 deficiency in *E. coli*. (**A**) Strain construction and genotype confirmation by PCR for *trmD-deg* and *trmD-cont* strains. Colony-PCR using primers targeting the 5’-terminal and 3’-flanking regions of *trmD* (shown by arrows) confirmed clones of *trmD-deg* and *trmD-cont* strains. The deg-tag adds to the C-terminus of TrmD with the YALAA peptide, which is targeted by the ClpXP degradation system, whereas the cont-tag does not. The clones were streak-purified until the WT band was completely removed. The mutant genomic *trmD* locus was introduced by P1 transduction to G78 cells, which lack the chromosomal *clpX* gene. (**B**) Growth of *trmD-deg* and *trmD-cont* G78 cells on an LB plate. An overnight culture of *trmD-deg* and *trmD-cont* cells in LB + Glc was serially diluted, spotted on an LB + Ara plate to turn on *clpXP* expression, and incubated at 37 °C overnight. (**C**) Representative growth of *trmD-deg* and *trmD-cont* G78 cells in a liquid LB culture. An overnight culture of *trmD-deg* and *trmD-cont* G78 cells in LB + Glc was grown in LB for 1-2 h, then diluted into LB + Ara at OD_600_ of 0.1 and grown to OD_600_ of 0.3 at 37 °C. The cycle of dilution and re-growth was repeated 3 times to observe a growth defect of the *trmD-deg* strain. (**D**) Western blot analysis of TrmD protein. An overnight culture of *trmD-deg* and *trmD-cont* cells in LB + Glc was diluted to LB + Ara at T = 0, and was sampled at the indicated time points. Cell lysates were separated on a 12% SDS-PAGE and TrmD protein was detected by primary antibody against *E. coli* TrmD and a secondary antibody against rabbit IgG. (**E**) Primer-extension inhibition analysis. An overnight culture of *trmD-deg* and *trmD-cont* cells in LB + Glc was diluted to LB + Ara at T = 0 and the fresh culture was taken through 3 cycles of dilution and re-growth. Total RNA was extracted and probed over the course of the 3 cycles of dilution and re-growth, each with a 5’-^32^P-labeled DNA primer targeting *E. coli* tRNA^Leu/CAG^. Primer extension by reverse transcription was analyzed on a 12% PAGE/7M urea gel and phosphor-imaging. Primer extension would terminate in TrmD deficiency at one nt downstream of m^1^G37, generating a 15-nt fragment, whereas primer extension would continue to the 5’-end in the control, generating a 53-nt fragment. The calculated fraction of m^1^G37 is shown for each sample. At the time of cell harvest, the m^1^G37 level for *trmD-deg* cells was below 25% at T = 4 h, whereas it was 100% for *trmD-cont* cells at T = 3 h.

Overnight cultures of *trmD-deg* and *trmD-cont* strains were grown in LB + Glc and were spotted onto an LB + Ara plate to turn on expression of *clpXP*. Analysis of a serial dilution of each strain confirmed that *trmD-deg* cells rapidly lost viability, whereas *trmD-cont* cells retained viability over several orders of dilution (*Figure 1B*). In liquid culture, in which each strain was freshly diluted into LB + Ara at OD_600_ 0.1 and grown to 0.3 - 0.4, followed by a second round of dilution and growth, we observed that the growth of the *trmD-deg* strain was retarded in the third round, whereas that of the *trmD-cont* strain was robust (*Figure 1C*). Although the growth defect of the *trmD-deg* strain only manifested in the third cycle of dilution, the level of TrmD protein drastically decreased in 30 min after the first dilution into fresh LB + Ara (*Figure 1C*), whereas that in the *trmD-cont* strain remained stable up to 90 min (*Figure 1D*) and longer (*Figure 1-figure supplement 1B*). We speculate that m^1^G37-methylated tRNAs, which were stable for hours even after TrmD had decreased (Masuda *et al*., 2019), allowed these cultures to grow for some time.

To measure cellular levels of m^1^G37 during this series of dilution and growth, we designed a 5’-^32^P-labeled primer of 14 nucleotides (14-nt) complementary to *E. coli* tRNA^Leu^(CAG). The presence of m^1^G37 would inhibit primer extension, producing a 15-nt product, whereas the absence of m^1^G37 would permit primer extension to the 5’-end, producing a 53-nt product. Analysis of total RNA samples, collected after the first dilution into fresh LB + Ara in the same time course (*Figure 1C*), showed that the 15-nt product progressively decreased in *trmD-deg* cells, indicating gradual loss of m^1^G37, but that it remained stable in *trmD-cont* cells (*Figure 1E*). The level of m^1^G37 was calculated as the fraction of the 15-nt product in the sum of the 15-nt and 53-nt products, whereas the level of the primer was not calculated due to its use in excess. At the time of cell harvesting, m^1^G37 in *trmD-deg* cells was at or below 25% (T = 4.2 h), whereas that in *trmD-cont* cells was 100% (T = 3 h). Collectively, these results demonstrate the ability of the degron approach to control the stability of TrmD and to produce conditional m^1^G37 deficiency in *E. coli* cells.

### Codon-specific ribosome stalling in m^1^G37 deficiency

We performed ribosome profiling and RNA-seq analyses on two biological replicates of the *trmD-deg* and *trmD-cont* strains, which were cultured in the presence of *clpXP* expression with three cycles of dilution and growth and were harvested at OD_600_ of 0.3. We also obtained a third set of samples using *trmD-KO (trm5–)* and *trmD-WT (trm5–)* strains, which were cultured in the absence of *trm5* expression in three cycles of dilution and growth and were harvested at OD_600_ of 0.3. Given that m^1^G37 is associated with a specific set of *E. coli* tRNAs, we looked in the ribosome profiling data for local differences in the rates of elongation for each codon in m^1^G37 deficiency. If loss of m^1^G37 impaired tRNA decoding, we expected that the ribosome would linger on affected codons to accumulate higher levels of density. To quantify these codon-specific effects, we defined a pause score for each codon as the ribosome density at the first nt of the codon normalized by the average ribosome density on the gene where that codon occurs. We computed the pause score for all of the 61 sense codons by averaging the pause score at thousands of instances of each codon across the entire transcriptome. Strikingly, we observed significant increases in the pause score for a set of codons when each was positioned at the ribosomal A site during decoding (*Figure 2*). Most notably, the pause score at Pro codons was dramatically increased in m^1^G37 deficiency, showing an increase from 1.4 in *trmD-cont* cells to 3.5 in *trmD-deg* cells on a log_2_ scale (x-axis and y-axis) (*Figure 2A*, left). Similarly, the pause score at Pro codons increased from 1.2 in *trmD-WT (trm5–)* cells to 2.5 in *trmD-KO (trm5–)* cells (*Figure 2A*, right). The increase in the pause score for all Pro codons indicates that m^1^G37 deficiency affected decoding by some or all of the isoacceptors of tRNA^Pro^, leading to strong ribosome pausing during elongation.

**Figure 2.**
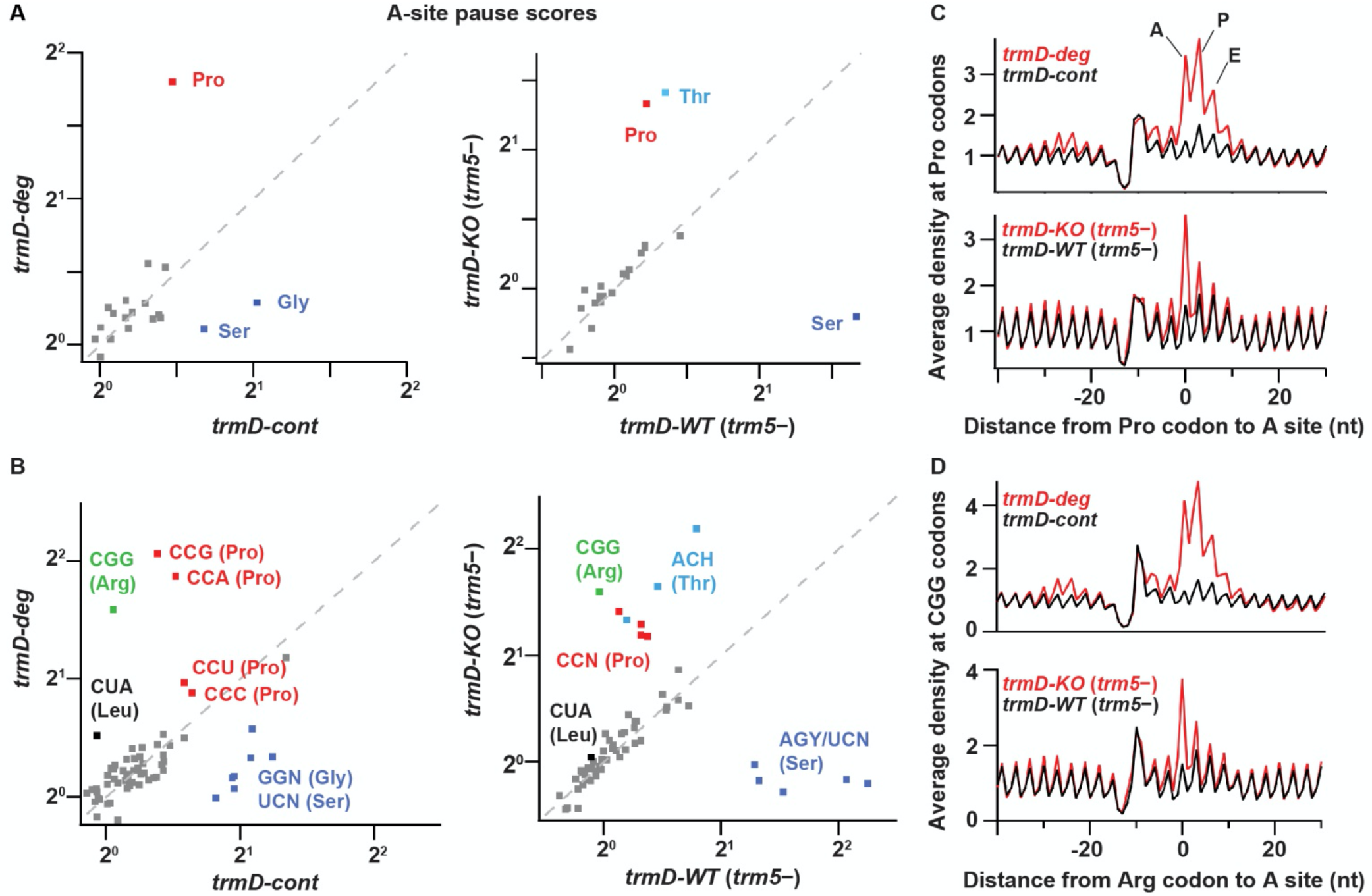
TrmD depletion leads to ribosome pausing at CCN (Pro) and CGG (Arg) codons. (**A**) Pause scores for codons positioned in the ribosomal A site. The codons are grouped by amino acid. Left panel: data from *trmD-deg* and *trmD-cont* strains upon ClpXP induction. Right panel: data from *trmD-KO* (*trm5–*) and *trmD-WT* (*trm5–*) strains upon turning off *trm5*. (**B**) The same as above but showing the 61 sense codons individually. (**C**) Plots of average ribosome density aligned at Pro codons for *trmD-deg* strain (top) and *trmD-KO* (*trm5–*) strain (bottom) in red with their respective control strains in black. The peaks corresponding to the ribosomal A, P, and E sites are labeled. (**D**) Plots of average ribosome density aligned at CGG codons as above. Note that *trmD-deg* samples were obtained with the traditional ribosome profiling lysis buffer whereas *trmD-KO* (*trm5–*) samples were obtained with a high Mg^2+^ lysis buffer that more effectively arrests translation after cell lysis, revealing these pauses at higher resolution.

We also observed differences in ribosome pausing at other codons (*Figure 2A*), which likely resulted from artifacts of the ribosome profiling method and not from translation differences due to m^1^G37 deficiency. There were stronger pauses at Ser and Gly codons in the control samples of *trmD-cont* than in the *trmD-deg* samples, and stronger pauses at Ser codons in the control samples of *trmD-WT (trm5–)* than in the *trmD-KO (trm5–)* samples. These pauses were likely due to artifacts of harvesting bacterial cultures. We previously observed pauses at Ser and Gly codons in cells harvested by filtration and showed that filtration reduced aminoacylation of the tRNA cognate to these codons (Mohammad *et al*, 2019). These Ser and Gly pauses, however, are less obvious in strains where protein synthesis is defective (e.g. due to loss of the elongation factor EFP) (Woolstenhulme *et al*, 2015), which may explain their lower pause scores in the *trmD-deg* and *trmD-KO (trm5–)* samples than in the control samples. We have also observed variable pausing at Thr codons under different harvesting conditions in unpublished work, although those pauses have not been as well characterized. Given that all of the samples described here were harvested by filtration, we attributed the pauses at Ser, Gly, and Thr codons (labeled in blue in *Figure 2A*) to artifacts of cell growth and harvesting and did not consider them further.

At the codon level, we observed higher pause scores for all four Pro codons (CCN, in red, *Figure 2B*) in m^1^G37-deficient cells relative to control cells, as shown in *trmD-deg* vs. *trmD-cont* strains (left) and in *trmD-KO (trm5–)* vs*. trmD-WT (trm5–)* strains (right). In both replicates of the *trmD-deg* strain, Pro codons CCG and CCA had significantly higher pause scores than CCC and CCU. In contrast, all four CCN codons showed similar pause scores in the *trmD-KO (trm5–)* strain. While the reason for these differences between the two strains is not clear, the implication is that ribosomes paused at all four Pro codons in both strains. Also, we observed elevated pause scores on the CGG codon (in green) in m^1^G37 deficiency, increasing from 1.0 in control cells to 3.0 in *trmD-deg* cells and in *trmD-KO (trm5–)* cells. Notably, CGG was the only Arg codon where pausing was observed upon loss of m^1^G37, whereas the other Arg codons CG[A/C/U], AGA, and AGG were not affected. Additionally, we observed a small increase of the pause score at the Leu codon CUA in m^1^G37 deficiency at 1.4 relative to the pause score of 1.0 in control samples (*Figure 2B*). Together, these data showed significant pauses at all Pro codons CCN, the Arg CGG codon, and to a lesser extent the Leu CUA codon. All of the paused codons are translated by tRNAs that are substrates of m^1^G37 methylation, although pauses at Leu codons CU[C/G/U], which are translated by tRNAs that should also be methylated with m^1^G37, were not observed.

The increase in the average pause scores in m^1^G37 deficiency indicates accumulation of ribosome density when the affected codon is positioned in the A site. To explore the possibility of other effects on translation, we looked more broadly at the average ribosome density aligned to a specific codon of interest. For *trmD-deg* vs. *trmD-cont* samples, we observed strong increases of ribosome density in the A, P, and E sites at Pro codons, and at the Arg CGG codons in m^1^G37 deficiency, with the highest peak in the ribosomal P site (*Figure 2C-D*, top). Although these findings could be interpreted to mean that m^1^G37 deficiency in tRNAs for these codons led to pausing when each tRNA was in the P or E site, we recognized that the *trmD-deg* and *trmD-cont* samples were obtained using the traditional lysis buffer with chloramphenicol to arrest translation in the lysate. We have since discovered that ribosomes continue to translate a few codons in the lysate under these conditions (Mohammad *et al*., 2019), suggesting that the several peaks observed in m^1^G37 deficiency likely arose from a strong pause at the A site that was blurred by ongoing translation in the lysate during sample preparation. In contrast, the *trmD-KO (trm5–)* and *trmD-WT (trm5–)* samples were prepared with a buffer containing 150 mM MgCl_2_, which immediately arrests ribosomes without the need for antibiotics (Mohammad *et al*., 2019). In these samples, we observed that pauses at Pro codons and at Arg CGG codons (*Figure 2C-D*, bottom) were the strongest when each codon was positioned at the A site, without an increased density in the P and E sites. We conclude that pauses are primarily in the A site, due to defects in decoding associated with m^1^G37 deficiency.

Interestingly, we observed increased ribosome density ∼25 nt upstream of affected codons in *trmD-deg* relative to *trmD-cont* samples (*Figure 2C-D*, top). This distance is roughly equivalent to the footprint length of a single ribosome, indicating that the increased density was due to collision of an upstream ribosome with a paused ribosome that was struggling to decode an affected codon. The collision of two ribosomes suggests that the pausing of the downstream ribosome at the affected codon is significantly prolonged. These findings are consistent with our prior observation of pausing and formation of disomes upon treatment of cells with mupirocin, an antibiotic that blocks aminoacylation of tRNA^Ile^ (Mohammad *et al*., 2019).

### Reduced aminoacylation and A-site peptide-bond formation of m^1^G37-deficient tRNAs

The observed ribosome pausing at specific codons in m^1^G37 deficiency raised two possibilities for the tRNAs translating these codons. First, m^1^G37 deficiency may reduce aminoacylation of these tRNAs by the respective aminoacyl-tRNA synthetases (aaRSs), preventing them from forming a ternary complex (TC) with EF-Tu-GTP and entering the ribosomal A site. Second, m^1^G37 deficiency may prevent these tRNAs from serving as efficient acceptors for peptidyl transfer, leading to inhibition of peptide-bond formation. We addressed these two possibilities, which are not mutually exclusive, with all three isoacceptors of tRNA^Pro^ and the tRNA^Arg^(CCG) isoacceptor. To isolate the effect of m^1^G37, we prepared tRNAs as T7 RNA polymerase transcripts lacking m^1^G37 or any other post-transcriptional modification (the G37-state) and compared their activity to transcripts that were subsequently modified with m^1^G37 by TrmD *in vitro* (the m^1^G37-state). For some of these tRNAs, we also purified from cells the native-state tRNA containing the full complement of natural post-transcriptional modifications.

Aminoacylation of tRNA^Pro^ with Pro was performed with purified *E. coli* ProRS under steady-state multi-turnover conditions (Zhang *et al*, 2006). The initial rate as a function of the concentration of each tRNA was measured and the data were fit to the Michaelis-Menten equation to derive kinetic parameters *K*_m_ (tRNA) and *k*_cat_. For all three isoacceptors of tRNA^Pro^, the catalytic efficiency *k*_cat_/*K*_m_ was decreased from the m^1^G37-sate to the G37-state by 3 to 10-fold (*Figure 3A and Figure 3–table supplement 1A*). The reduction in *k*_cat_/*K*_m_ was driven by an increase in *K*_m_ for all three tRNAs, indicating that m^1^G37 may be important for binding of each tRNA to ProRS (*Figure 3 – table supplement 1A*). This apparent binding defect makes sense structurally because m^1^G37 is immdediately downstream of the two conserved anticodon nucleotides G35-G36, the major determinants of tRNA^Pro^ binding to ProRS (Cusack *et al*, 1998; Yaremchuk *et al*, 2000; Yaremchuk *et al*, 2001).

**Figure 3.**
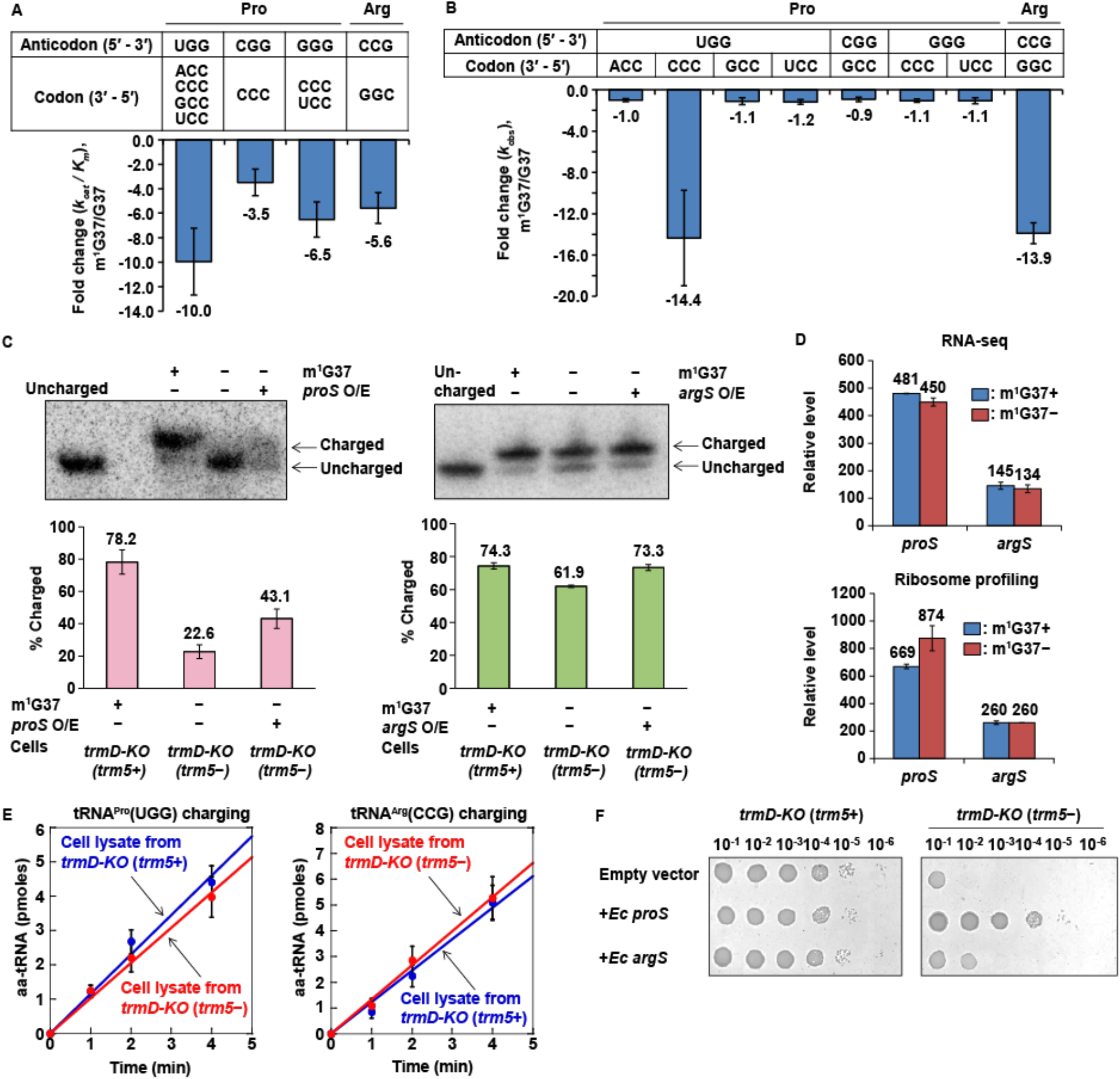
Loss of m^1^G37 reduces aminoacylation and peptide-bond formation of tRNAs at the A site. (**A**) Fold-change in the loss of *k*_cat_/*K*_m_ of aminoacylation of tRNAs from the m^1^G37-state to the G37-state. Bar graphs are shown for the UGG, the CGG, and the GGG isoacceptors of *E. coli* tRNA^Pro^, and for the CCG isoacceptor of *E. coli* tRNA^Arg^, with the fold-change value displayed at the bottom of each bar. The bars are SD of three independent (n = 3) experiments, and the data are presented as the mean ± SD for each sample. (**B**) Fold-change in the loss of *k*_obs_ of peptide-bond formation of tRNAs from the m^1^G37-state to the G37 state at the A site. Bar graphs are shown for the UGG, CGG, and GGG isoacceptors of *E. coli* tRNA^Pro^, and for the CCG isoacceptor of *E. coli* tRNA^Arg^, with pairing to the cognate codon, and the fold-change value is displayed at the bottom of each bar as the mean ± SD of three independent experiments (n = 3). (**C**) Northern blots of acid-urea gels showing the fractional distribution of charged vs. uncharged *E. coli* tRNA^Pro^(UGG) (left) and tRNA^Arg^(CCG) (right) under m^1^G37+ (+) and m^1^G37– (–) conditions, and over-expression (O/E) in m^1^G37– (–) condition of *proS* and *argS*, respectively. The uncharged tRNA^Pro^(UGG) and tRNA^Arg^(CCG) was run in parallel for each tRNA. The % charged tRNA species in each case is shown in the bar graph below (n = 3). (**D**) Relative levels of expression of *E. coli proS* and *argS* in RNA-seq and ribosome-profiling analysis in m^1^G37+ and m^1^G37– conditions. (**E**) Analysis of protein level of *proS* (left) and *argS* (right) in m^1^G37+ and m^1^G37– conditions by assaying for the aminoacylation activity in cell lysates against *E. coli* tRNA^Pro^(UGG) and tRNA^Arg^(CCG) in the m^1^G37-state (n = 3). (**F**) Viability of *E. coli* cells *trmD-KO* (*trm5+)* and *trmD-KO (trm5–)* cells with an empty vector, or the vector over-expressing *E. coli proS* or *argS*.

The largest decrease in *k*_cat_/*K*_m_ (10-fold) upon loss of m^1^G37 was observed for the UGG isoacceptor. Unique among the isoacceptors of tRNA^Pro^, the UGG isoacceptor is required for cell growth and survival (Nasvall *et al*, 2004) and is most critically dependent on m^1^G37 for maintaining the translational reading-frame (Gamper *et al*., 2015a). The critical role that m^1^G37 plays in the UGG isoacceptor was further highlighted by the finding that aminoacylation reaction with the m^1^G37-state tRNA have the same *k*_cat_/*K*_m_ values as the native-state tRNA (*Figure 3 – table supplement 1A*), indicating that all other post-transcriptional modifications played little or no role in aminoacylation.

Aminoacylation of the tRNA^Arg^(CCG) isoacceptor with Arg was performed with purified *E. coli* ArgRS under steady-state multi-turnover conditions. The *k*_cat_/*K*_m_ of aminoacylation was decreased from the m^1^G37-state to the G37-sate by 5.6-fold. (*Figure 3A and Figure 3–table supplement 1A*), and this decrease was largely driven by a loss in *k*_cat_ (*Figure 3–table supplement 1A*), indicating that m^1^G37 contributed to catalysis. This is a different effect than what we observed for tRNA^Pro^, where loss of m^1^G37 reduced tRNA binding to ProRS. Additionally, we observed a loss of 14.3-fold in *k*_cat_/*K*_m_ from the native-state to the G37-state (*Figure 3–table supplement 1A*), greater than the 5.6-fold loss due to m^1^G37 alone, indicating that other post-transcriptional modifications played a role in aminoacylation of this tRNA^Arg^, which also contrasts the observation that m^1^G37 alone is sufficient for rapid aminoacylation of tRNA^Pro^. Together, these results show that m^1^G37 is required for efficient aminoacylation of all three isoacceptors of tRNA^Pro^ and the tRNA^Arg^(CCG) isoacceptor, and that loss of m^1^G37 reduces aminoacylation in all cases, although by apparently different mechanisms.

To determine whether m^1^G37 deficiency affected peptidyl transfer to affected tRNAs in the A site, we used our *E. coli in vitro* translation system of purified components and supplemented it with requisite tRNAs and translation factors to perform a series of ensemble rapid kinetic studies (Gamper *et al*., 2021; Gamper *et al*., 2015a; Gamper *et al*, 2015b). We assayed for the synthesis of a peptide bond between the [^35^S]-fMet moiety of the P-site [^35^S]-fMet-tRNA^fMet^ of a 70S initiation complex (70S IC) and the aminoacyl moiety of a TC delivered to the A site. This assay monitors all of the reactions at the A site, including the initial binding of the TC to the A site, EF-Tu-catalyzed GTP hydrolysis, accommodation of the aa-tRNA to the A site, and peptidyl transfer. Each TC carried a fully aminoacylated tRNA in the G37-state or in the m^1^G37-state, in excess of the 70S IC, to allow evaluation of the activity of peptide-bond formation. The 70S IC was kept at a limiting concentration relative to the *K*_d_ of peptide-bond formation (Ledoux *et al*, 2009), such that the measured *k*_obs_ represented the catalytic efficiency *k*_cat_/*K*_m_ of peptidyl transfer. For tRNA^Pro^, the UGG isoacceptor was assayed at all four Pro codons CCN, the CGG isoacceptor was assayed at the codon CCG, and the GGG isoacceptor was assayed at the codons CC[C/U] (*Figure 3B and Figure 3–table supplement 1B*). The results of these assays showed that, of all of the tested anticodon-codon pairs, loss of m^1^G37 only had a significant effect on *k*_obs_ with the UGG isoacceptor at the CCC codon, decreasing *k*_obs_ from the m^1^G37-state to the G37-state by 14.4-fold (*Figure 3B and Figure 3–table supplement 1B*). Thus, in contrast to aminoacylation, where all isoacceptors of tRNA^Pro^ were affected by loss of m^1^G37, loss of m^1^G37 only affected peptidyl transfer for one isoacceptor at one codon. Likewise, the CCG isoacceptor of tRNA^Arg^ was assayed at the cognate codon CGG, showing a significant decrease of *k*_obs_ by 13.9-fold from the m^1^G37-state to the G37-state (*Figure 3B and Figure 3–table supplement 1B*). Thus, for the single isoacceptor of tRNA^Arg^, loss of m^1^G37 reduced the activity of both aminoacylation and peptide-bond formation.

Following up on the results of these kinetic studies *in vitro*, we investigated whether m^1^G37 deficiency led to the loss of aminoacylation of TrmD-dependent tRNAs *in vivo*. Total RNA was extracted from *E. coli trmD-KO (trm5+)* and *trmD-KO (trm5–)* cells using an acid buffer (pH 4.5) and run on an acid-urea gel to preserve the levels of charged aa-tRNA vs. uncharged tRNA at the time of harvest. Northern blots with probes against the tRNA^Pro^(UGG) and tRNA^Arg^(CCG) isoacceptors showed that both tRNAs had reduced aa-tRNA levels, supporting the notion that m^1^G37 deficiency decreased aminoacylation *in vivo* (*Figure 3C*). Compared to the drastic reduction in aminoacylation levels of Pro-tRNA^Pro^(UGG) (78 to 22%), the reduction for Arg-tRNA^Arg^(CCG) was milder (74 to 62%) (*Figure 3C*), indicating that aminoacylation of tRNA^Pro^(UGG) was much more sensitive to m^1^G37 deficiency in cells. RNA-seq and ribosome profiling data confirm that the lower aminoacylation levels were not due to loss of expression of the corresponding *proS* or *argS* genes (*Figure 3D*). Moreover, cell lysates in m^1^G37-abundant (*trm5*+) and m^1^G37-deficient (*trm5*–) conditions exhibited a similar aminoacylation activity when assayed with a m^1^G37-tRNA substrate, either tRNA^Pro^(UGG) or tRNA^Arg^(CCG) (*Figure 3E*), indicating that the enzymatic activity of *proS* and *argS* was similar between the two growth conditions.

Conversely, over-expression of *proS* and *argS* in *trmD-KO* (*trm5*–) cells increased the aminoacylation level of tRNA^Pro^(UGG) and tRNA^Arg^(CCG), respectively (*Figure 3C*), indicating that the poor aminoacylation kinetics of m^1^G37-deficient tRNA can be overcome by increasing the corresponding aaRS enzyme. We asked whether these higher levels of aminoacylation would improve the translation activity of m^1^G37-deficient tRNAs and restore cell viability. Indeed, *trmD-KO (trm5+)* cells grew robustly, regardless of the expression level of *proS* or *argS*, whereas *trmD-KO (trm5–)* cells that normally grew much more poorly showed increased viability upon over-expression of the respective gene (*Figure 3F*). The recovery of cell viability was stronger upon over-expression of *proS* than *argS*, arguing that low levels of aminoacylation of the tRNA^Pro^ species was the driver limited cell growth under m^1^G37 deficiency.

Taken together, these findings show that m^1^G37 deficiency reduces the aminoacylation level of all tRNA^Pro^ species and tRNA^Arg^(CCG) *in vitro* and *in vivo*; this loss of aminoacylation levels compromises cell viability but can be overcome by over-expression of *proS* and to a lesser extent over-expression of *argS*.

### Changes in gene expression by m^1^G37 deficiency

We next asked what changes in gene expression took place in m^1^G37 deficiency. Using the DESeq algorithm to analyze the RNA-seq data, we identified genes whose steady-state levels of mRNAs were altered with statistical significance by loss of m^1^G37. We found 220 genes with more than 2-fold higher expression (*p* < 0.01) in the *trmD-deg* strain (shown in red, *Figure 4A*). Conversely, we identified 166 genes that were repressed by more than 2-fold (*p* < 0.01, colored in blue, *Figure 4A*). For both the up- and down-regulated genes, we identified several pathways that are affected and whose genes are enriched at statistically significant levels (*Figure 4C*). We also observed changes at the translational level. Using the Xtail algorithm to identify changes in ribosome occupancy (RO = Ribo-seq density / RNA-seq density), we found 71 genes with higher levels of apparent ribosome occupancy (1.7 to 6.5-fold increase) in the *trmD-deg* strain relative to the *trmD-cont* strain (*p* < 0.01, *Figure 4B*). Because the magnitude of transcriptional changes was greater than that of translational changes, yielding clearer results on which pathways are affected, we focused on the transcriptional changes arising from m^1^G37 deficiency.

**Figure 4.**
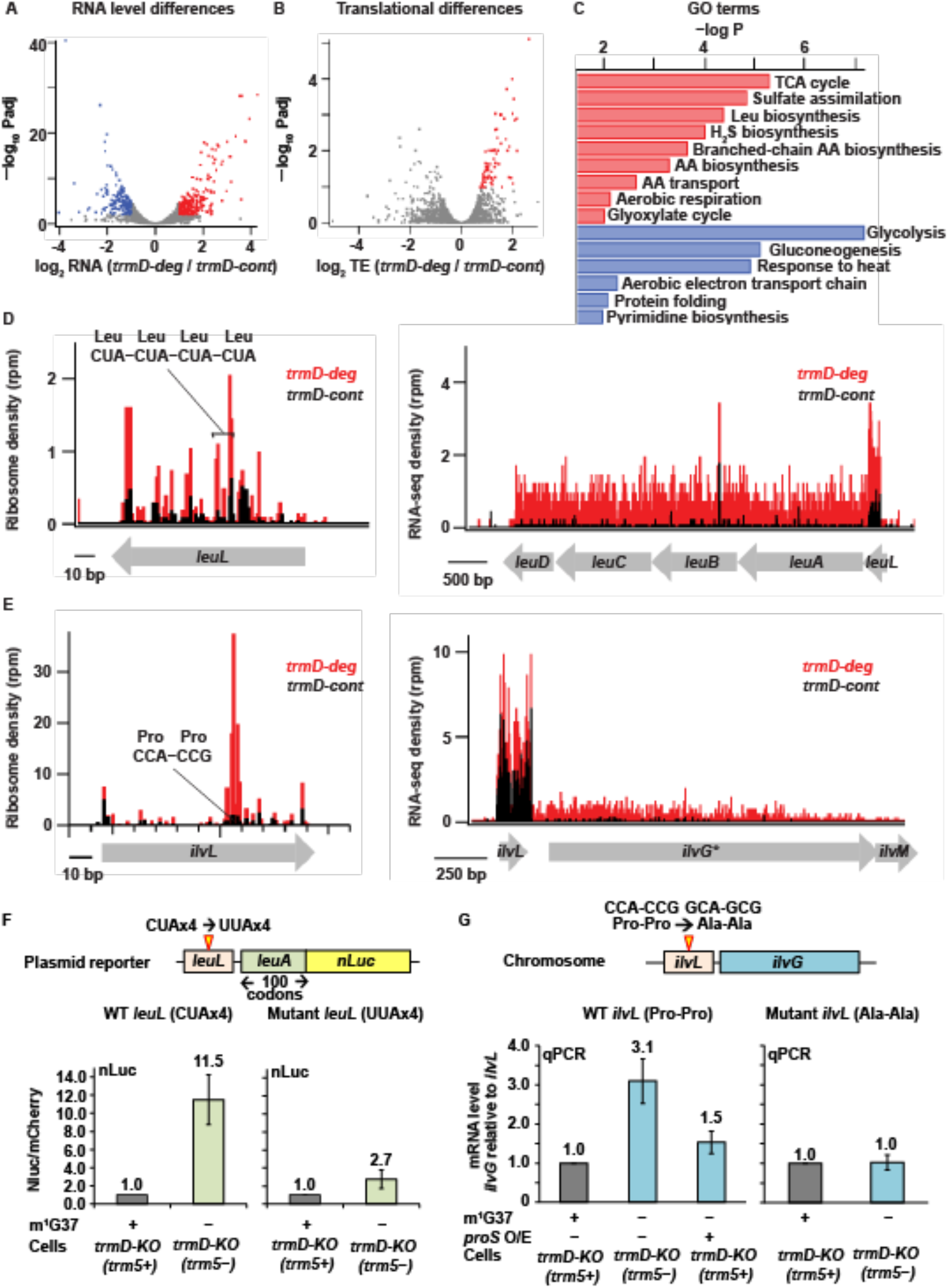
Changes in gene expression arising from m^1^G37 deficiency. (**A**) Volcano plot showing differences in steady state RNA levels (from RNA-seq data analyzed by DESeq) using samples of two replicates of *trmD-deg* and *trmD-cont* strains upon ClpXP induction. Genes that are more than 2-fold higher in the mutant with *p* < 0.01 are colored red (n = 220); those that are 2-fold lower with *p* < 0.01 are colored blue (n = 166). (**B**) Volcano plot showing differences in ribosome occupancy (Ribo-seq / RNA-seq analyzed by Xtail) using samples from two replicates of *trmD-deg* and *trmD-cont* strains upon ClpXP induction. Of these, 95 genes that are up-regulated (p < 0.1) are colored red. (**C**) Gene ontology categories for genes enriched in the up-regulated genes (red) and down-regulated genes (blue) from (A). (**D**) Gene model showing ribosome reads (in rpm) and RNA-seq density (in rpm, reads per million mapped reads) at the *leuL* leader sequence upstream of the *leuABCD* operon. (**E**) Gene model showing ribosome reads (in rpm) and RNA-seq density (in rpm) at the *ilvL* leader sequence upstream of the *ilvGMEDA* operon. (**F**) A plasmid reporter construct that demonstrates the codon-specific effect in the *leuL* leader sequence on expression of the downstream *leuA* gene. The plasmid reporter encodes the genomic sequence of *leuL,* followed by the first 100 codons of *leuA*, in the native sequence or with substitution of the four consecutive CUA codons with four UUA codons. The 100 codons of *leuA* are then fused to the nLuc gene in-frame. Expression of nLuc, normalized by co-expression of mCherry in a separate plasmid within the same cell, is shown for m^1^G37+ and m^1^G37– conditions (n = 3). (**G**) Analysis of the CCA-CCG (Pro-Pro) codons in the genomic locus of *ilvL* as the determinant of regulation of gene expression of the downstream *ilvG* gene. A variant *E. coli* strain was created that changed the CCA-CCG (Pro-Pro) codons to GCA-GCG (Ala-Ala) codons at the natural genomic locus, using CRISPR/Cas editing. Expression of *ilvG* was monitored by qPCR analysis in m^1^G37+ and m^1^G37– conditions and in m^1^G37– condition with over-expression of *proS* (n = 3).

Many of the genes with higher RNA levels upon TrmD depletion are involved in amino acid biosynthesis and transport (*Figure 4C*). For example, induction of the Leu operon (*leuA*, *leuB*, *leuC*, and *leuD*) was particularly strong (∼16-fold). This Leu operon is regulated by transcriptional attenuation depending on the efficiency of synthesis of the upstream *leuL* leader peptide (Wessler & Calvo, 1981; Wohlgemuth *et al*, 2013). As with leader sequences of other amino acid biosynthesis operons (Kolter & Yanofsky, 1982), translation of *leuL* serves as a sensitive genetic switch that has evolved to sense levels of aa-tRNAs and to respond to amino acid starvation by up-regulating the downstream biosynthetic pathways (Wohlgemuth *et al*., 2013). We observed ribosomal pausing in *leuL* at the four consecutive Leu CUA codons in m^1^G37 deficiency (*Figure 4D*), consistent with our observation of ribosome pausing at CUA codons (*Figure 2B*), suggesting that the pausing would prevent transcriptional termination and allow transcription elongation into the downstream Leu biosynthetic genes, resulting in elevated RNA levels. In agreement with this transcriptional attenuation mechanism, we observed in the control *trmD-cont* strain efficient transcriptional termination downstream of *leuL*, where the average RNA-seq density (adjusted for length) decreased 29-fold from the *leuL* leader sequence to the first gene in the operon (*leuA*), whereas we observed in the *trmD-deg* strain only a 4-fold decrease (*Figure 4D*). These results support the notion that ribosome pausing at the four consecutive Leu CUA codons in *leuL* during m^1^G37 deficiency reduced transcriptional termination of the downstream *leuA* gene (*Figure 4D*). Another example is the *ilvL* leader sequence upstream of the *ilvGMEDA* operon encoding genes for biosynthesis of branched chain amino acids Ile, Leu, and Val. Ribosome pausing in *ilvL* would prevent transcriptional termination and allow expression of the downstream biosynthetic genes (Nargang *et al*, 1980). Although this genetic switch has evolved to sense changes in levels of aa-tRNAs associated with the amino acids produced by the downstream genes, we instead observed strong ribosome pauses at two consecutive Pro codons CCA-CCG in m^1^G37 deficiency (*Figure 4E*). These strong pauses within the *ilvL* leader sequence could trigger anti-termination, explaining the higher RNA-seq levels of the downstream gene *ilvG*. Indeed, while we observed in the control strain a 45-fold decrease in the RNA-seq density from the region around *ilvL* to the first half of the downstream *ilvG* gene, we observed in the *trmD-deg* strain only a 9-fold decrease in the RNA-seq density (*Figure 4E*). Notably, the *ilvG* gene in strains derived from *E. coli* K12 MG1655, such as the one used here, has a mutation that induces a frameshift midway through the gene, suggesting that it may be a pseudogene, although the *ilvG* gene appears intact in other *E. coli* strains.

We confirmed that the changes of expression of these two operons were due to pauses at specific codons induced by m^1^G37 deficiency in cognate tRNAs. To examine the effect of the *leuL* leader sequence on expression of *leuA*, we generated a plasmid reporter construct containing *leuL* and the first 100 codons of *leuA* fused in-frame to the nano-lucifierase (nLuc) gene *(Figure 4F*). Upon expression of the WT reporter construct, m^1^G37 deficiency in *trmD-KO (trm5–)* cells elevated the nLuc readout by 11.5-fold relative to control cells, consistent with the notion that ribosome stalling at the four consecutive Leu CUA codons in *leuL* prevents transcriptional termination and allows transcription of the dowstream *leuA-nLuc* fusion. In contrast, a smaller activation during m^1^G37 deficiency (only 2.7-fold) was observed with a second nLuc reporter where the four consecutive Leu CUA codons were changed to Leu UUA codons, which do not require m^1^G37 for translation.

To examine the effect of the *ilvL* leader sequence on expression of *ilvG*, we altered the two Pro codons CCA-CCG in *ilvL* to two Ala codons GCA-GCG by genome editing using CRISPR and followed the expression of the downstream *ilvG* gene with qPCR (*Figure 4G*). With the wild-type *ilvL* sequence, *ilvG* was induced 3.1-fold in m^1^G37-deficient *trmD-KO (trm5–)* cells relative to m^1^G37-abundant *trmD-KO (trm5+)* cells (left panel, bar 2). In contrast, no induction was observed for the mutant *ilvL* sequence (right panel, bar 2). These results support the notion that ribosome stalling at the Pro codons in *ilvL* activates expression of the downstream *ilvG* in m^1^G37 deficiency. Furthermore, given our observation that overexpression of *proS* restored cell viability under m^1^G37 deficiency (*Figure 3F*), we asked if it might also minimize pausing and induction of this genetic switch. We found that over-expression of *proS* in *trmD-KO (trm5–)* cells with the wild-type *ilvL* leader sequence reduced the activation of *ilvG* from 3.1-fold to 1.5-fold (left panel, bars 2 and 3). This result is consistent with the notion that elevated *proS* expression restores aminoacylation levels of tRNA^Pro^, even in m^1^G37 deficiency, relieving ribosome stalling and repressing the downstream *ilvG* gene.

In addition to Leu, Ile, and Val, the biosynthetic pathways for Trp, His, and Cys were highly up-regulated in the *trmD-deg* strain (*Figure 4C*), even though there is no evidence of ribosome stalling at the 5’-leader of the relevant operons. The up-regulation of these pathways suggests that they are activated by a genome-wide response due to ribosome stalling at Pro CCN, Arg CGG, and Leu CUA codons (*Figure 2B*). The increase in expression of the Cys biosynthesis pathway is correlated with the enrichment in GO terms for genes involved in sulfate assimilation and hydrogen sulfide biosynthesis (*Figure 4C*) and is consistent with the notion that cysteine is synthesized from serine in bacteria by incorporation of sulfide to *O*-acetylserine (Kredich & Tomkins, 1966).

### Changes of gene expression in m^1^G37 deficiency consistent with the stringent response

The high levels of expression of amino acid biosynthesis genes in m^1^G37 deficiency was reminiscent of the bacterial stringent response, which is triggered in nutrient starvation by uncharged tRNAs binding to the ribosomal A site, activating synthesis of ppGpp catalyzed by the RelA protein (Gourse *et al*., 2018). In the stringent response, ppGpp binds to two sites on RNA polymerase (RNAP) and re-programs the transcriptional landscape (Gourse *et al*., 2018; Ross *et al*, 2016; Ross *et al*, 2013), shutting down gene expression of rRNAs and ribosome proteins, while up-regulating amino acid biosynthesis genes to respond to amino acid starvation. Our observation that m^1^G37 deficiency reduced aminoacylation of all isoacceptors of tRNA^Pro^ and the tRNA^Arg^(CCG) isoacceptor raised the possibility that these effects would induce the programmatic changes in gene expression similar to those in the stringent response.

To test this possibility, we compared the changes in RNA levels in m^1^G37 deficiency with the changes in RNA levels upon induction of the stringent response. We used the dataset for the stringent response obtained by Gourse and co-workers that compared changes in RNA levels between an *E. coli* strain containing the ppGpp-binding sites in RNAP and a variant lacking the binding sites upon over-expression of a *relA* mutant that was deleted of a regulatory domain and was constitutively activated to synthesize ppGpp (Sanchez-Vazquez *et al*, 2019). This dataset focused on *E. coli* genes that exhibited transcriptional up- and down-regulation in response to direct binding of ppGpp to RNAP, rather than indirect effects from starvation-induced changes in metabolism. After 5 minutes of induction of the *relA* mutant, the dataset identified a set of 401 down-regulated genes (by > 2-fold), a set of 321 up-regulated genes (by > 2-fold), and a set of 3,036 genes with less than 2-fold changes (Sanchez-Vazquez *et al*., 2019). Using these same sets of genes, we found in our RNA-seq data that genes that were down-regulated upon *relA* over-expression were also down-regulated in m^1^G37 deficiency (log_2_(median) = –0.36; *t*-test *p* = 8.9 × 10^−15^) (*Figure 5A*). Likewise, genes that were up-regulated in *relA* over-expression were also up-regulated in m^1^G37 deficiency (*Figure 5A*), although the extent of up-regulation in m^1^G37 deficiency was smaller (log_2_(median) = 0.29; *t*-test *p* = 2.3 × 10^−11^). Additionally, the non-responsive genes showing less than 2-fold changes upon *relA* over-expression also showed little change in m^1^G37 deficiency (log_2_(median) ∼ 0.04). Together, these results show that genes that are strongly induced or repressed by the stringent response are affected in a similar way by m^1^G37 deficiency, arguing that m^1^G37 deficiency induces changes in gene expression similar to those of the stringent response.

**Figure 5.**
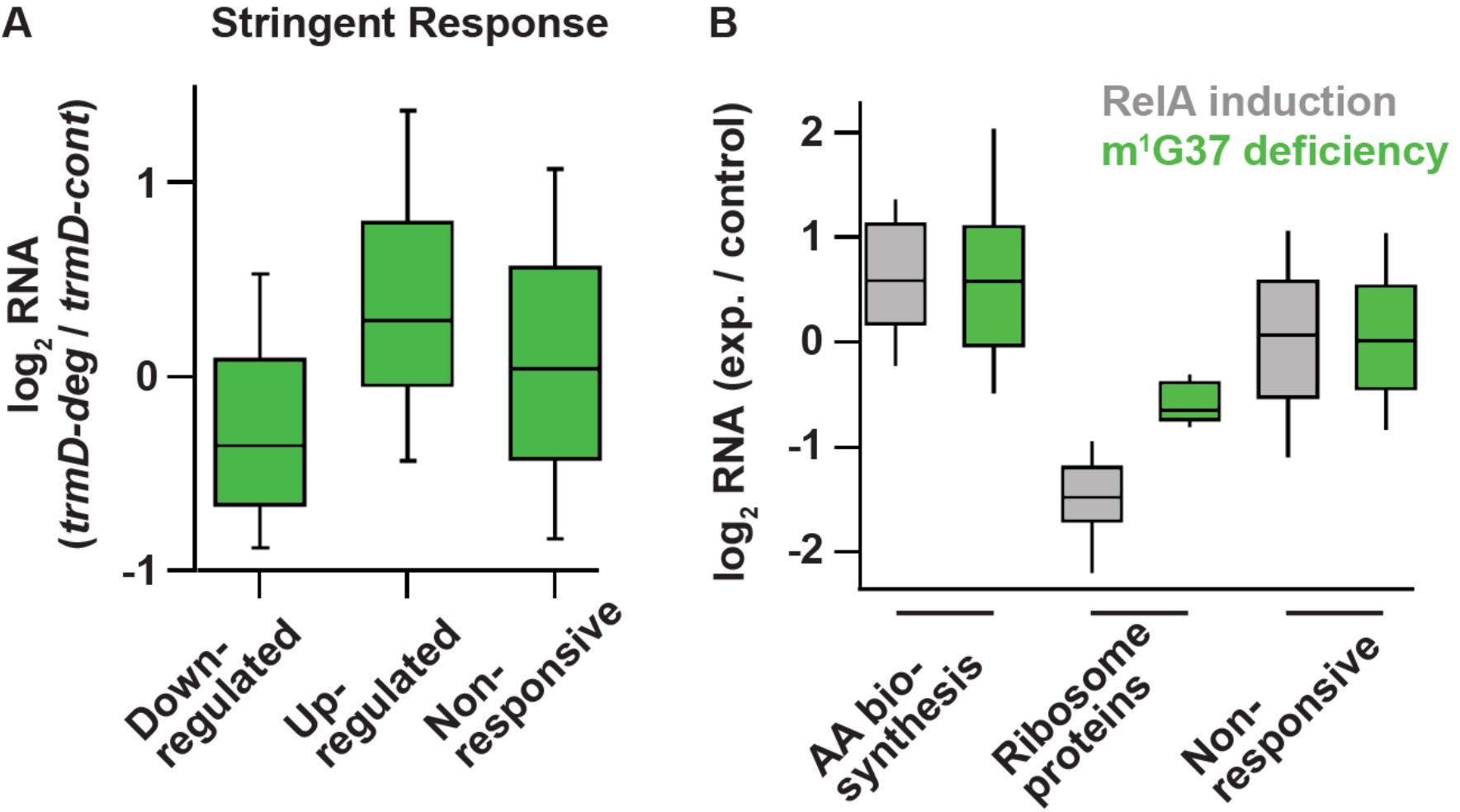
Deficiency of m^1^G37 induces gene expression similar to that of the bacterial stringent response. (**A**) A set of genes published previously by Gourse et al. (Sanchez-Vazquez *et al*., 2019), whose steady state RNA levels changed more than 2-fold upon induction of the stringent response by RelA over-expression (down n = 401 genes, up n = 321 genes, other = 3036 genes). Using these same sets of genes, the ratio of RNA levels in *trmD-deg* and *trmD-cont* strains upon ClpXP induction are shown. (**B**) Changes in expression in genes involved in amino-acid biosynthesis (n = 89), genes encoding ribosomal proteins (n = 47), and other genes.

Following up on this observation, we next asked to what extent two characteristic pathways known to be regulated by the stringent response were also affected by m^1^G37 deficiency. In nutrient starvation, bacterial cells down-regulate ribosome biosynthesis due to the reduced demand for protein production, but up-regulate amino acid biosynthesis (Gourse *et al*., 2018). As expected, the published RNA-seq data upon *relA* over-expression showed that 89 genes associated with amino acid biosynthesis in the EcoCyc database were more highly expressed than other genes (log_2_(median) = 0.59, Mann-Whitney *p* = 4.8 × 10^−12^). Intriguingly, we found that these genes were also up-regulated to the same extent in m^1^G37 deficiency (log_2_(median) = 0.58, Mann-Whitney *p* = 1.2 × 10^−8^) (*Figure 5B*), consistent with the enrichment of these genes in our GO annotation (*Figure 4C*). Likewise, 47 ribosome protein genes were down-regulated upon *relA* over-expression (log_2_(median) = –1.48, Mann-Whitney *p* = 1.6 × 10^−-24^) and they were also down-regulated in m^1^G37 deficiency, although to a lesser extent (log_2_(median) = –0.64, Mann-Whitney *p* = 1.1 × 10^−12^) (*Figure 5B*). As a control, we also found that the genes that were not induced in these sets (i.e., not ribosome proteins and not involved in amino-acid biosynthesis) were less responsive to *relA* over-expression, and that they were also less responsive in m^1^G37 deficiency (*Figure 5B*). Collectively, the parallel changes in steady-state RNA levels between m^1^G37 deficiency and *relA* over-expression were remarkable, suggesting that loss of m^1^G37 triggers a response similar to that of the *relA*-dependent stringent response.

### Metabolic changes

Some of the most striking changes in gene expression in m^1^G37 deficiency occurred in central metabolic pathways: glycolysis, the citric acid (the tricarboxylic (TCA)) cycle, and fatty-acid oxidation. For example, ten genes associated with the TCA cycle were significantly up-regulated in the *trmD-deg* strain, making this pathway the most enriched in the GO terms (*Figure 4C*). We identified genes of the TCA cycle that showed log_2_-fold changes of expression in the two samples of *trmD-deg* replicates (*Figure 6A*, column 1 and 2 from the left) and in the two samples from the stringent response study in the ppGpp-binding sites in RNAP (column 3) or without the binding sites (column 4), respectively (Sanchez-Vazquez *et al*., 2019). For the top half of the cycle, there are strong increases in expression in the *trmD-deg* strains for genes responsible for taking malate to oxaloacetate (*mqo*), condensing oxaloacetate with acetyl-CoA to form citrate (*gltA),* and isomerizing citrate to isocitrate (*acnAB*). In contrast, for the lower half of the cycle, including genes for the two decarboxylation reactions from isocitrate to 2-oxoglutarate and then to succinate, there was little or no significant change. It appears that, rather than a simple up-regulation of the entire TCA cycle, genes for the specific steps that are shared with the glyoxylate cycle were increased.

**Figure 6.**
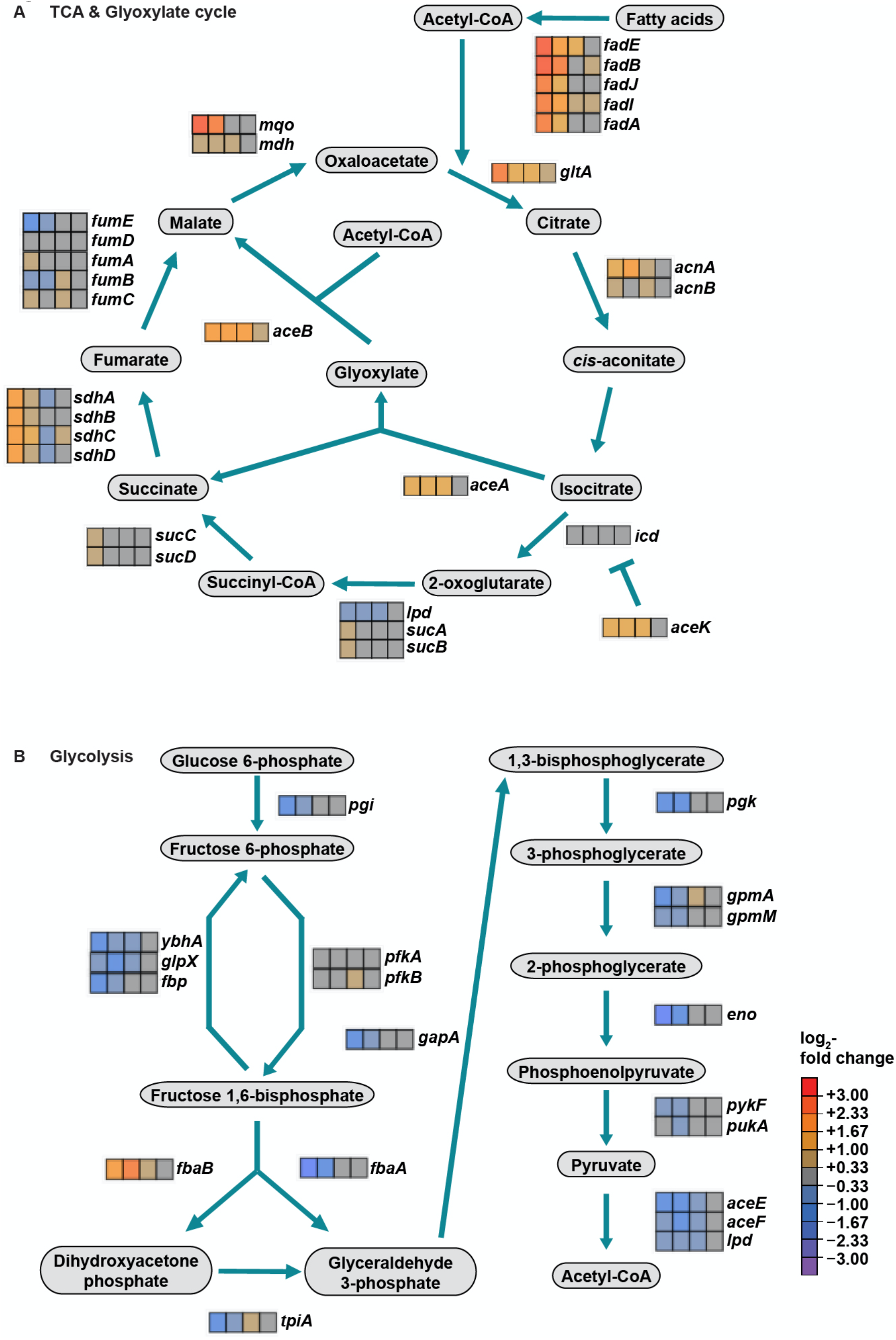
Changes in central metabolic pathways in m^1^G37 deficiency. (**A**) Depiction of the TCA cycle and the glyoxylate cycle and its associated genes showing the log_2_-fold change in four RNA-seq samples. From left to right, these samples are: two replicates of the *trmD-deg* strain (obtained in this study) and two samples where constitutively active *relA* was overexpressed to induce the stringent response, first in a strain with the wild-type RNA polymerase, and second in a strain with two mutations that render RNA polymerase insensitive to ppGpp (Sanchez-Vazquez *et al*., 2019). (**B**) Depiction of the glycolysis pathway and associated genes and the same four RNA-seq samples with the same heat map scale (shown in (**A**).

The glyoxylate cycle bypasses the oxidative decarboxylation steps in the TCA cycle and directly converts isocitrate into malate and succinate. The purpose of these bypass reactions is to replenish TCA cycle intermediates, using the two-carbon acetyl-CoA subunits to build up metabolic intermediates rather than generating energy through decarboxylation. The key enzymes in the glyoxylate cycle are isocitrate lyase (encoded by *aceA*) and malate synthase (encoded by *aceB*), whose genes were up-regulated in the *trmD-deg* strain by 2.6- and 3.2-fold, respectively. In order to direct isocitrate through these enzymes in the glyoxylate cycle rather than through isocitrate dehydrogenase (encoded by *icd*) and the standard TCA cycle, the *aceK* kinase phosphorylates and inactivates isocitrate dehydrogenase. As with *aceA* and *aceB*, we found that *aceK* is also highly up-regulated in m^1^G37 deficiency. Up-regulation of the glyoxylate cycle is likely an anabolic alternative to reserve carbon resources for utilization in amino acid biosynthesis. Notably, *aceA*, *aceB*, and *aceK* are also up-regulated by *relA* over-expression in a manner dependent on ppGpp binding to RNAP, suggesting that the glyoxylate cycle may play a role in amino acid biosynthesis during the canonical stringent response.

Analysis of the 166 genes that were repressed by more than 2-fold (*Figure 4A*) underscored the metabolic changes taking place during m^1^G37 deficiency. The pathway with the strongest representation of GO terms was glycolysis, where 10 genes associated with glycolysis were repressed in m^1^G37 deficiency (e.g. 4-fold down at the RNA level for the gene encoding enolase). The down-regulation of these genes was not due to losses in the available sugars in the media between the *trmD-deg* and *trmD-cont* strains, both of which were cultured and grown in the same way. Down-regulation of glycolysis is consistent with the activation of the glyoxylate cycle, which is typically repressed during metabolism of glucose but is activated during metabolism of two-carbon compounds such as acetyl-CoA. Rather than arising from catabolism of glucose, the acetyl-CoA arises from fatty-acid oxidation whose enzymes are highly up-regulated in m^1^G37 deficiency. It seems that m^1^G37 deficiency moves metabolism away from consuming glucose towards consuming fatty acids. These changes were only seen in the *trmD-deg* replicates, but not upon *relA* over-expression and induction of the stringent response, possibly due to the much longer period of growth in the *trmD-deg* samples (several hours) compared to the short 5 min induction of *relA* in the stringent response study. Alternatively, loss of TrmD may induce additional changes above and beyond those associated with the canonical stringent response.

## Discussion

Here we provide a genome-wide view of the effect of m^1^G37 deficiency on protein synthesis upon depletion of TrmD in *E. coli*. This genome-wide view is important, because TrmD is ranked as a high-priority antibacterial target (White & Kell, 2004), due to its fundamental distinction from Trm5, conservation throughout the bacterial domain, essentiality for bacterial life, and possession of a small molecule-binding site for drug targeting. The small molecule-binding site in TrmD is unique, consisting of a protein topological knot-fold that holds the methyl donor *S*-adenosyl methionine in an unusual shape (Ahn *et al*, 2003; Elkins *et al*, 2003; Ito *et al*, 2015), which is different from that in Trm5 and in most other methyl transferases (Goto-Ito *et al*, 2009), indicating the possibility to explore novel chemical space and diversity of drugs targeting TrmD. Additionally, we have shown that TrmD deficiency can disrupt the double-membrane cell-envelope structure of Gram-negative bacteria, thus facilitating permeability of multiple drugs into cells, preventing efflux of these drugs, and accelerating bactericidal action (Masuda *et al*., 2019). This multitude of advantages of targeting TrmD emphasizes that a better understanding of the genome-wide function of its m^1^G37 product will help develop a successful antibacterial strategy.

Using ribosome profiling, we show that m^1^G37 deficiency causes ribosome stalling at codons specific for tRNAs that are methylated by TrmD, including strong pauses at all Pro CCN codons and the Arg CGG codon and weak pauses at the Leu CUA codon. Stalling is primarily observed when the affected codons are in the ribosomal A site, indicating a distinct defect than that of +1 frameshifting, which occurs in m^1^G37 deficiency after decoding at the A site and during tRNA translocation to the P site and occupancy within the P site (Gamper *et al*., 2021; Gamper *et al*., 2015a). The importance of m^1^G37 at the ribosomal A site thus expands the spectrum of its biology, which can be summarized in a model that explains its indispensability throughout the entire elongation cycle of protein synthesis (*Figure 7*). In this model, m^1^G37 deficiency induces ribosome stalling in the A site, triggering changes in gene expression and possibly affecting mRNA stability and co-translational protein folding. Ribosome stalling can be prolonged if a subset of the affected tRNAs is unable to perform peptide-bond formation in the A site, leading to ribosome collisions that increase the frequency of frameshifting (Smith *et al*, 2019). Even if some of the m^1^G37-deficient tRNAs manage to enter the A site and participate in peptide-bond formation, they would induce +1 frameshifting during translocation to the P site and destabilize interaction with the ribosomal P site (Hoffer *et al*., 2020), leading to premature fall-off from the ribosome and termination of protein synthesis. In contrast to starvation that can be alleviated by re-supply of nutrients, this series of defects associated with m^1^G37 deficiency cannot be readily resolved, thus underscoring the essentiality of m^1^G37 in protein synthesis that is coupled to cell growth and survival.

**Figure 7.**
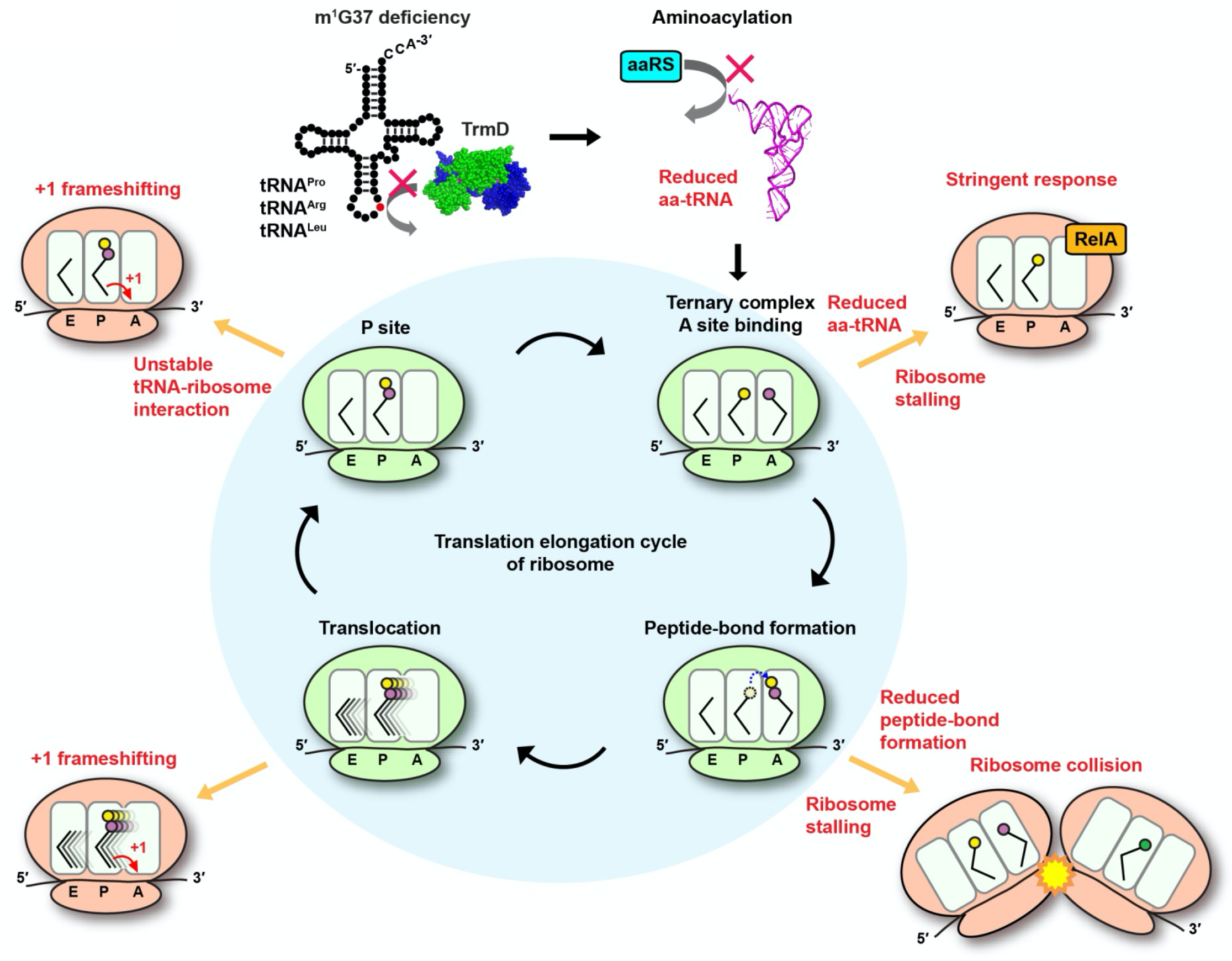
Deficiency of m^1^G37 impairs the elongation cycle of protein synthesis in bacteria. A cloverleaf structure of tRNA, showing the position of m^1^G37 in red. Loss of m^1^G37 reduces aminoacylation, lowering levels of aa-tRNAs at the ribosome A site, stalling the ribosome, and activating programmatic changes in gene expression similar to those induced by the bacterial stringent response upon RelA sensing of uncharged tRNA and synthesis of ppGpp. Ribosome stalling at the A site is prolonged if the m^1^G37-deficient aa-tRNA is delayed in peptide-bond formation, leading to ribosome collision. Even if the m^1^G37-deficient tRNA manages to enter the A site participate in peptide-bond formation, it would promote +1 frameshifting during translocation from the A site to the P site and during occupancy within the P site, while also destabilizing the P site structure and possibly falling off from the P site to prematurely terminate protein synthesis. Accumulation of these defects ultimately collapses the entire elongation cycle and leads to cell death.

An important finding of this work is that ribosome stalling in m^1^G37 deficiency is primarily driven by loss of aminoacylation of affected tRNAs, which is shown in enzyme-based kinetic assays for all isoacceptors examined, and in cell-based acid-urea assays for tRNA^Pro^(UGG) and tRNA^Arg^(CCG), two major TrmD-dependent species in bacteria. While the loss of aminoacylation in m^1^G37 deficiency is not due to the loss of ProRS or ArgRS in cells, it can be restored by over-expression of either enzyme, which improves cell viability. These results clearly demonstrate that the loss of aminoacylation in m^1^G37 deficiency, and consequently the accumulation of uncharged tRNAs, is the underlying basis of ribosome stalling that induces significant changes in gene expression. Direct changes include up-regulation of the *leuABCD* and *ilvGMEDA* operons. Indirect changes include hundreds of genes that are activated similar to those in the bacterial stringent response – the central bacterial response to nutrient starvation upon RelA sensing of uncharged tRNAs binding to the A site of translating ribosomes. These changes certainly make sense given that we see high levels of deacylated tRNAs accumulating in m^1^G37 deficiency and that the binding of deacylated tRNA to the A site is known to induce (p)ppGpp synthesis by RelA on the ribosome. We did not test the role that RelA likely plays in the indirect changes in gene expression in m^1^G37 deficiency, because the *trmD-deg* and *trmD-KO (trm5–)* cells are very sick and the deletion of *relA* is expected to further reduce their viability. Further confounding the analysis is that *E. coli* contains two genes responsible for ppGpp synthesis: *relA* and *spoT*, which have overlapping functions. Upon *relA* deletion, the protein product of *spoT* can still synthesize ppGpp in response to some of the metabolic changes that are observed here. Nonetheless, the changes in gene expression in *trmD* deficiency largely parallel with those observed during the stringent response, consistent with the notion that the loss of aminoacylation in m^1^G37 deficiency activates the stringent response.

Previous studies have shown that loss of the metabolic state of tRNA can induce changes in gene expression similar to those induced by cellular response to nutrient starvation. Examples include the loss of the 5-methoxycarbonylmethyl (mcm^5^) and/or the 2-thio (s^2^) groups from the normal tRNA in the mcm^5^s^2^U34-state (Nedialkova & Leidel, 2015; Zinshteyn & Gilbert, 2013), loss of the threonyl-carbamoyl group from the normal t^6^A37-state (Thiaville *et al*, 2016), and loss of the inosine (I) group from the normal I34-state (Lyu *et al*, 2020). Among these, the lack of the s^2^ group from the mcm^5^s^2^U34-state is known to reduce the binding and accommodation of tRNA to the ribosomal A site and the rate of translocation to the P site (Ranjan & Rodnina, 2017). In contrast, while m^1^G37 deficiency reduces peptide bond formation for some tRNAs at the A site, it consistently reduces the rate of aminoacylation for all tRNAs examined, which has not been shown for other metabolically deficient tRNAs. The rate reduction in aminoacylation of tRNA^Pro^ is uniformly manifested at the *K*_m_ (tRNA) step but not the *k*_cat_ step (*Figure 3 – supplement Table 1*), which may be a feature of the bacterial ProRS that discriminates against m^1^G37-deficient tRNA due to the proximity of the methylation to the adjacent anticodon G36 nucleotide that is recognized by the bacterial enzyme. In contrast, the human ProRS has diverged in recognition of the tRNA anticodon by shifting emphasis toward the G35 nucleotide, which is more distant from m^1^G37 (Burke *et al*, 2001). This shift raises the possibility that the human ProRS may better accommodate m^1^G37-deficient tRNAs for aminoacylation, and that the eukaryotic response to m^1^G37 deficiency is executed in a step other than aminoacylation during decoding at the A site. This possibility remains to be addressed in future studies.

The molecular insight obtained from this study is important for antibacterial therapeutics. It demonstrates that m^1^G37 is required for the ribosomal activity at the A site and, when combined with previous studies, it expands the importance of m^1^G37 to the entire elongation cycle of protein synthesis. This suggests that drug targeting of TrmD can be used in combination with classic antibiotics that target the A site (e.g., streptomycin, paromomycin), the translocation reaction (e.g., chloramphenicol, sparsomycin), and the P site (e.g., macrolides) (Arenz & Wilson, 2016). Harnessing this new insight will enable developing novel and improved strategies to combat the rapid emergence of multi-drug resistance of bacterial pathogens.

## Materials and methods

### Construction of *trmD-deg* and *trmD-cont* strains

*E. coli trmD-deg* strain was constructed following the published protocol (Carr *et al*., 2012). The degron tag was amplified from the template DNA (provided by Dr. Sean Moore), which encodes a FLAG tag, a His_6_ tag, and the degron sequence YALAA followed by the promoter and the coding sequence of the tetracycline resistance gene. This region was amplified with primers homologous to the 3’-end and flanking sequence of the chromosomal *trmD* locus. The PCR product was purified and electroporated into competent cells of the recombinogenic *E. coli* strain SM1405 expressing λRed recombinase (Datsenko & Wanner, 2000). Recombination was confirmed by PCR analysis of colonies on plates containing tetracycline using primers homologous to the 3’-end of the chromosomal *trmD* locus. The P1 lysate of the confirmed *trmD-deg* strain was used for transduction into the recipient *E. coli* G78 strain whose chromosomal *clpX* was deleted (Carr *et al*., 2012). A resultant P1 transductant was selected on plates with tetracycline, confirmed by PCR, and transformed with a library of the plasmid p*clpPX* harboring a cassette encoding *clpP* and *clpX* and random mutations at the promoter region that control expression of the cassette (Carr *et al*., 2012). Transformants were screened on LB + Ara (0.2%) plates to turn on expression of the cassette and to identify the clone with the highest efficiency of the degron activity as indicated by Western blots. *E. coli trmD-cont* strain was created in the same procedure, except that the degron tag coding for the YALAA sequence was placed after the stop codon of *trmD*. The P1 transductant harboring *trmD-cont* in the G78 strain was transformed with the p*clpPX* plasmid that showed the highest degron activity, generating the *trmD-cont* control strain. The primer sequences are shown below:

> Forward recombination: CGGAACACGCACAACAGCAACATAAACATGATGGGATGGCGGGTGGCTCCGACTACAAGG
>
> Reverse recombination: ATAATTTAATCTCTTATCCTGGGTAAACTGATATCTCGGGGGCTTAGGTCGAGGTGGCCC
>
> Forward confirmation at the *trmD* locus: ATGTGGATTGGCATAATTAGCCTGTTTCC
>
> Reverse confirmation at the *trmD* locus: GAATTCCGGTTACGAATAGCGATAACCACGCC

### Growth conditions

*trmD-deg* and *trmD-cont* cells were grown in LB + Glc at 37 °C overnight, inoculated into 20 mL of fresh LB + Ara at 1:100, and grown for 2 h at 37 °C into the start of the log phase. This growth cycle was repeated three times. In the first cycle, cells were inoculated into 20 mL of LB + Ara (0.2%) and 10 mM Ser at OD_600_ of 0.1 and grown for 1 h at 37 °C to OD_600_ of ∼0.4, at which point TrmD protein was no longer detectable by Western blots. In the second cycle, cells were inoculated into 100 mL of LB + Ara (0.2%) and 10 mM Ser at OD_600_ of 0.1 and grown for 1 h at 37 °C to OD_600_ of ∼0.4. In the third cycle, cells were inoculated into 500 mL at OD_600_ of 0.1 and grown for 2-3 h at 37 °C to OD_600_ of ∼0.3. Cells from 200-300 mL culture were harvested by rapid filtration (see below). *trmD*-*KO (trm5–)* and *trmD*-*WT (trm5–)* cells were grown in LB + Ara (0.2%) at 37 °C overnight and were then taken through three cycles of growth in MOPS Medium (EZ Rich Defined, Teknova) containing Glc as the only carbon source. In the first cycle, cells in the overnight culture were inoculated into 10 mL of MOPS at OD_600_ of 0.1 and grown for 4 h at 37 °C to turn off the Ara-dependent expression of *trm5* and to deplete m^1^G37-tRNAs. In the second cycle, cells were inoculated into 25 mL of MOPS + Glc and grown for 3 h, and in the third cycle, cells were inoculated into 300 mL of MOPS + Glc and grown 2-4 h at 37 °C until OD_600_ of ∼0.3. Cells from 200-300 mL culture were harvested by rapid filtration (see below).

### Western blots

*E. coli* G78 strains of *trmD-deg* and *trmD-cont* mutants were grown overnight in LB (no Ara) and were diluted 1:100 into fresh LB + Ara (0.2%). Cells were grown at 37 °C and were lysed 0, 15, 30, 45, 60, and 90 min after inoculation. Cell lysate of 15 μg of protein was separated on 12% SDS-PAGE and levels of TrmD were probed by the primary anti-TrmD antibody (generously provided by Dr. Glenn Bjork) at a dilution of 1:10,000 and a secondary anti-rabbit IgG antibody (Sigma-Aldrich) at a dilution of 1:160,000. Signals were detected by the SuperSignal West Pico Chemiluminescent Substrate (Thermo Fisher Scientific in the Chemi-Doc XRS+ System (Bio-Rad).

### Primer-extension inhibition assays

Primer-extension inhibition analyses of *E. coli* tRNA^Leu/CAG^ were performed as described (Masuda *et al*., 2019) on *trmD-deg* and *trmD-cont* lysates at the indicated time points after switching to LB + Ara. A DNA primer targeting the sequence of nucleotides 40 to 54 of the tRNA was chemically synthesized, ^32^P-labeled at the 5’-end by T4 polynucleotide kinase, annealed to the tRNA, and extended by Superscript III reverse transcriptase (Invitrogen) at 200 U/µL with 6 μM each dNTP in 50 mM Tris-HCl, pH 8.3, 3 mM MgCl_2_, 75 mM KCl, and 1 mM DTT at 55 °C for 30 min. The reaction was heated at 70 oC for 15 min and quenched with 10 mM EDTA. cDNA products were separated by 12% PAGE/7M urea and visualized by phosphorimaging. In these assays, the extension product of the read-through cDNA is 53-nt in length, whereas the extension inhibition product is 15-nt in length.

### Cell harvesting and lysis

We used two different cell harvesting strategies to arrest ribosomes and block translation after cell lysis. The strategy of cell harvesting can influence the quality of ribosome profiling data (Mohammad *et al*., 2019). *trmD-deg* and *trmD-cont* cultures were harvested by rapid filtration on a Kontes 99 mm filtration apparatus and 0.45 µm nitrocellulose filter (Whatman) and the cells were flash-frozen in liquid nitrogen. Cells were lysed in lysis buffer (20 mM Tris-HCl, pH 8.0, 10 mM MgCl_2_, 100 mM NH_4_Cl, 5 mM CaCl_2_, 100 U/mL RNase-free DNase I, and 1 mM chloramphenicol). Due to sequence-specific inhibition by chloramphenicol (Marks *et al*, 2016; Mohammad *et al*, 2016; Nakahigashi *et al*, 2014; Orelle *et al*, 2013), this cell harvesting strategy was more prone to artefacts at the codon level, creating apparent pauses when the smaller amino acids Ala, Gly, and Ser were at the penultimate position in nascent polypeptide chains. Furthermore, chloramphenicol does not fully arrest translation in cell lysates, blurring the signal (Mohammad *et al*., 2019). In contrast, *trmD-KO (trm5–)* and *trmD-WT (trm5–)* cultures, while also harvested by rapid filtration and flash-freezing, were lysed in a buffer lacking chloramphenicol but containing high concentrations of MgCl_2_ (20 mM Tris-HCl, pH 8.0, 150 mM MgCl_2_, 100 mM NH_4_Cl, 5 mM CaCl_2_, 0.4% triton X100, 0.1% NP-40, and 100 U/mL RNase-free DNase I). In this lysis buffer, the high MgCl_2_ inhibits translation by preventing ribosomal conformational changes required for elongation (Mohammad *et al*., 2019). After lysis, the samples in the high MgCl_2_ buffer were centrifuged over a sucrose cushion to collect polysomes, which were resuspended in the standard lysis buffer without the high salt to allow micrococcal nuclease (MNase) digestion and isolation of ribosome-protected mRNA fragments for deep sequencing analysis.

### Library preparation

A 10-54% sucrose density gradient was prepared in the Gradient Master 108 (Biocomp) in the gradient buffer (20 mM Tris-HCl, pH 8.0, 10 mM MgCl_2_, 100 mM NH_4_Cl, and 2 mM DTT). A cell lysate of 5-20 AU was loaded to the sucrose density gradient and centrifuged in a SW41 rotor at 35,000 rpm for 2.5 h at 4 °C. Fractionation was performed on a Piston Gradient Fractionator (Biocomp). Libraries for ribosome profiling and RNA-seq were prepared as described (Mohammad *et al*., 2016; Woolstenhulme *et al*., 2015) on RNA fragments 15-45 nt in length, with the exception of RNA-seq libraries for the *trmD-KO (trm5–)* and *trmD-WT (trm5–)* samples, which were prepared using the TruSeq Stranded Total RNA kit (Illumina). The libraries were analyzed on a high-sensitivity BioAnalyzer (Agilent), and sequenced on the HiSeq2500 Illumina instrument.

### Analysis of sequencing data

The adaptor sequence CTGTAGGCACCATCAATAGATCGGAAGAGCACACGTCTGAA-CTCCAGTCA was removed by Skewer version 0.2.2 (Jiang *et al*, 2014). After reads mapping to tRNA and rRNA were removed, the remaining reads were aligned to *E. coli* MG1655 genome NC_000913.2 using bowtie version 1.1.2, requiring unique mapping sites and allowing two mismatches (Langmead *et al*, 2009). The position of the ribosome was assigned using the 3’-end of reads. Reads 10-40 nt in length were included in analyses unless otherwise specified for all samples except for RNA-seq samples from *trmD-KO (trm5–)* and *trmD-WT (trm5–)* prepared using the TruSeq kit, which were 50 nt in length.

For calculating pause scores, only reads 24-40 nt in length were used in *trmD-KO (trm5–)* and *trmD-WT (trm5–)* ribosome profiling samples because there was uncertainty about the position of the ribosome on reads shorter than 24 nt due to the different lysis buffer. In calculating pause scores, we only included genes with more than 0.1 reads per codon on average. The first seven and last seven codons in each ORF were ignored. For each instance of a codon of interest, we calculated a pause score by taking the density at the first nt only in the A site codon and dividing it by the mean density on the ORF. The average pause scores reported (*Figure 2*) represent the mean of the scores for all instances (typically thousands) of the codon of interest.

For analyses of changes in gene expression, we calculated for each gene the density of the RNA-seq or Ribo-seq in units of RPKM and then used the DESeq and Xtail packages in R with two replicates of the *trmD-deg* and *trmD-cont* samples to compute the log_2_ fold change in expression and the –log(P_adj_) value shown in *Figure 4*. Enrichment for GO terms were determined using the functional annotation tools in DAVID at david.ncifcrf.gov (Huang *et al*, 2007). RNA-seq data were mapped onto metabolic pathways using the Pathway Collage tool at ecocyc.org (Karp *et al*, 2021; Keseler *et al*, 2017).

### Aminoacylation of tRNA *in vitro*

Each tRNA was aminoacylated with the cognate amino acid by the respective aaRS enzyme that had been over-expressed in BL21 (DE3) and purified via binding to a Ni-NTA resin and eluted from the column by imidazole (Zhang *et al*., 2006). Each tRNA was heat-denatured at 85 °C for 3 min and re-annealed at 37 °C for 15 min. Aminoacylation in steady state conditions was performed at 37 °C in a 30 μL reaction of 0.25 - 20 μM tRNA, 5 nM aaRS, and 20 μM [^3^H]-amino acid (Perkin Elmer, 7.5 Ci/mmol) in a buffer containing 20 mM KCl, 10 mM MgCl_2_, 4 mM dithiothreitol (DTT), 0.2 mg/mL bovine serum albumin (BSA), 2 mM ATP (pH 8.0), and 50 mM Tris-HCl, pH 7.5 (Liu *et al*, 2011). Reaction aliquots of 5 µL were removed at different time intervals and precipitated with 5% (w/v) trichloroacetic acid on filter pads for 10 min twice. Filter pads were washed with 95% ethanol twice, with ether once, air dried, and measured for radioactivity in Tri-Carb 4910 TR scintillation counter (Beckman). Counts were converted to pmoles using the specific activity of the [^3^H]-amino acid after correcting for signal quenching by filter pads. Data corresponding to the initial rate of aminoacylation as a function of the tRNA substrate concentration were fit to the Michaelis-Menten equation to derive the *K*_m_ (tRNA), *k*_cat_ (catalytic turnover of the enzyme), and *k*_cat_/*K*_m_ (tRNA) (the catalytic efficiency of aminoacylation).

### Aminoacylation of tRNA in cell lysate

Harvested cells were resuspended in a lysis buffer (50 mM Tris-HCl pH 8.0 and 150 mM NaCl) and were disrupted by a Bioruptor Pico device (Diagenode) through 12 cycles of sonication and intermittent rest at 10 sec and 45 sec, respectively. The generated cell lysate was cleared of debris by ultracentrifugation through MTX-150 with the S100-AT4 rotor (Thermo Scientific) at 500,000 × *g* for 2 h at 4 °C, and was concentrated through Amicon Ultra-4 Centrifugal Filter Units (Millipore) to ∼0.5 mL. The concentrated lysate was washed with 4 mL of the binding buffer (10 mM Tris-HCl pH 7.5, 30 mM KCl, 10 mM MgCl_2_, and 1 mM DTT) and concentrated again through the same Amicon unit to 1.6 mL. Two aliquots of 300 µL DEAE sepharose were washed with 1 mL water by spinning at 3,000 × *g* for 2 min, and two washes with 1 mL of the binding buffer each. The 1.6 mL concentrated cell lysate was divided into two, and each was mixed with the washed DEAE sepharose and incubated at 4 °C with rotation for 1 h. The slurry was transferred to a gravity column and washed with 10 volumes of 1.2 mL of the binding buffer. The aaRS-containing fraction was eluted by 4 mL of the elution buffer (10 mM Tris-HCl pH 7.5, 400 mM KCl, 10 mM MgCl_2_, and 1 mM DTT), concentrated by two Amicon Ultra-0.5 mL Centrifugal Filters down to 100 µL, exchanged with the dialysis buffer (20 mM Tris-HCl pH 7.5, 130 mM KCl, 10 mM MgCl_2_, and 1 mM DTT), followed by dialysis in a Slide-A-Lyzer MINI Dialysis Device, 10k MWCO (Thermo Scientific) to remove small molecules. The post-DEAE fraction was determined for protein concentration by the Bradford assay. Molar concentration of proteins in the post-DEAE cell lysate was estimated based on an average molecular weight of *E. coli* proteins at 40 kDa.

Transcripts of *E. coli* tRNA^Pro^(UGG) and tRNA^Arg^(CCG) were made by *in vitro* transcription with T7 RNA polymerase and methylated with m^1^G37 by purified recombinant *E. coli* TrmD (Christian *et al*., 2004). Intracellular concentration of ProRS and ArgRS in the post-DEAE lysate was estimated assuming each representing 1% of total proteins to allow calculation of the molar concentration of each enzyme. To determine the intracellular aminoacylation activity of each enzyme, the post-DEAE lysate was incubated with a heat-cooled tRNA transcript at 37 °C in the aminoacylation buffer at the final concentration of 20 nM ProRS or ArgRS, 200 nM tRNA, and 20 mM KCl, 10 mL MgCl_2_, 4 mM DTT, 0.2 mg/mL BSA, 2 mM ATP, 50 mM HEPES pH 7.5, 20 µM [^3^H]-proline or [^3^H]-arginine (Lipman *et al*, 2000). Aliquots were removed at 1, 2, and 4 min, spotted onto a Whatman 3 MM filter pad, and precipitated by 5% trichloroacetic acid (TCA). Filters were washed with 5% TCA twice, 70% ethanol once, 100% ether once, and air-dried. The radioactivity measured by a Tri-Carb 4910 TR scintillation counter (Perkin-Elmer) was converted to pmoles of aa-tRNA synthesis.

To investigate the effect of over-expression of aaRSs, *E. coli trmD-*KO strain in BL21(DE3) maintained by human *trm5* (Gamper *et al*., *Nat Commun* 2015) was transformed with plasmid pET22b expressing *E. coli proS*, *argS*, or an empty vector. Overnight cultures in MOPS medium with 0.15% Ara + 0.05% Glc were inoculated 1:100 to fresh MOPS medium with 0.15% Ara + 0.05% Glc to induce the m^1^G37+ condition, or to MOPS with 0.2% Glc to induce the m^1^G37– condition, each harboring 0.025 mM isopropyl-β-D-thiogalactoside (IPTG) to induce expression of the plasmid-borne gene. After 2 cycles of growth and dilution at 37 °C, cells were harvested at OD_600_ 0.3-0.5 for isolation of total RNA and for preparation of cell lysate. To assay viability on a plate, overnight cultures in LB medium supplemented with 0.2% Ara were inoculated to fresh LB without Ara and grown for 3 h to deplete pre-existing Trm5 protein and m^1^G37-tRNAs. After depletion, cells were made by a 10-fold serial dilution, spotted on M9 plates containing 0.2% Ara or 0.2% Glc as the only carbon source, and grown at 37 °C for overnight. To test the effect of the Pro-Pro CCA-CCG codon motif on gene expression of *ilvL*, mutations were introduced to change the motif to Ala-Ala GCA-GCG on the *E. coli* chromosome of *trmD-*KO/MG1655 strain, using CRISPR-Cas9 mutagenesis (Reisch and Prather, *Sci Rep* 2018). Cells were grown and harvested for total RNA isolation for qPCR as above, but without addition of IPTG.

### Peptide-bond formation assays *in vitro*

Peptide-bond formation assays were performed on mRNA sequences that varied in the second codon position (shown as XXX below), while all other nucleotides (including the Shine-Dalgarno sequence (underlined) and the AUG start codon (bold face)) were the same.

> 5’-GGGA<u>AGGAGG</u>UAAAA**AUG**-XXX-CGU-UCU-AAG-(CAC)_7_-3’

Each reaction was carried out in 50 mM Tris-HCl, pH 7.5, 70 mM NH_4_Cl, 30 mM KCl, 3.5 mM MgCl_2_, 1 mM DTT, and 0.5 mM spermidine at 20 °C unless otherwise indicated (Gamper *et al*., 2015a; Liu *et al*., 2011). *E. coli* 70S ICs were formed by incubating 70S ribosomes, mRNA, [^35^S]-fMet-tRNA^fMet^, and initiation factors (IFs) 1, 2, and 3 with GTP for 25 min at 37 °C in the reaction buffer. Separately, each *E. coli* tRNA (4 μM) was aminoacylated with the cognate amino acid by the cognate aaRS (1 μM) to rapidly achieve plateau-level of aminoacylation as confirmed by enzymatic assays. To form a TC, the aminoacylation reaction was incubated with activated EF-Tu-GTP, which was formed separately for 15 min at 37 °C, and incubated in an ice bath for 15 min. To monitor peptide-bond formation, 70S ICs templated with each specified mRNA were mixed with a TC in an RQF-3 Kintek chemical quench apparatus. Final concentrations in these reactions were 0.1 µM for the 70S IC; 1.0 µM for mRNA; 0.5 µM each for IFs 1, 2, and 3; 0.25 µM for [^35^S]-fMet-tRNA^fMet^; 1.0 µM for EF-Tu for each aa-tRNA; 0.2 µM each for the aa-tRNAs; and 1 mM for GTP. Reactions were conducted at 20 °C unless otherwise specified, and were quenched by adding KOH to 0.5 M. After a brief incubation at 37 °C, aliquots of 1.8 µL were spotted onto a cellulose-backed plastic thin layer chromatography sheet and electrophoresed at 1000 V in PYRAC buffer (62 mM pyridine, 3.48 M acetic acid, pH 2.7) until the marker dye bromophenol blue reached the water-oil interface at the anode. The position of the origin was adjusted to maximize separation of the expected oligopeptide products. The separation of unreacted [^35^S]-fMet from each of the [^35^S]-fMet-peptide products was visualized by phosphorimaging and quantified using ImageQuant (GE Healthcare) and kinetic plots were fitted using Kaleidagraph (Synergy software).

### Acid-urea gel and Northern analysis

Total RNA was extracted from cells in an acidic condition (pH 4.5) to maintain the aminoacylated state of tRNAs as described (Parker *et al*., *Cell Systems* 2020). Cell cultures at the appropriate OD were mixed at a 1:1 volume ratio with 10% TCA and incubated on ice for 10 min. Cells were spun down at 8,000 rpm (= 7,600 × *g*) for 5 min at 4 °C, resuspended in the extraction buffer (0.3 M NaOAc, pH 4.5 and 10 mM EDTA), and mixed with an equal volume of ice-cold phenol-chloroform-isoamyl alcohol (83:17:3.4), pH 5.0. The mixture was vortexed for 5 cycles of 1 min vortex and 1 min rest on ice, and spun at 12,000 rpm (= 13,500 × *g*) for 10 min at 4 °C. The RNA-containing aqueous phase was mixed with an equal volume of cold isopropanol and incubated at −20 °C for 30 min for precipitation, followed by centrifugation at 14,000 rpm for 20 min at 4 °C to collect the pellet, which was washed with 70% ethanol and 30% 10 mM NaOAc, pH 4.5, and dried. The RNA-containing pellet was dissolved in a gel-loading buffer (10 mM NaOAc, pH 4.5 and 1 mM EDTA). Due to the high lability of the prolyl linkage in Pro-tRNA^Pro^ to hydrolysis (Peacock *et al*, 2014), total RNA was immediately prepared to run on an acid-urea gel. Assuming that tRNA accounts for 10% of total RNA, we made calculation to load 700 ng of total tRNA of each sample, mixed with 2 volumes of the acid-urea loading buffer (0.1 M NaOAc, pH 5.0, 9 M urea, 0.05% bromophenol blue (BPB), and 0.05% xylene cyanol (XC)). An intermediate size of acid-urea gel (14 × 17 cm) was used to resolve Pro-tRNA^Pro^ from uncharged tRNA^Pro^, whereas a Bio-Rad standard size mini gel was used to resolve Arg-tRNA^Arg^ from uncharged tRNA^Arg^. Each acid-urea gel was made with 6.5% polyacrylamide, 7 M urea, and 0.1 M NaOAc, pH 5.0, and run at 250 V for 3 h 45 min at 4 °C for the intermediate size gel and at 100 V for 2 h 30 min for the mini gel. Following electrophoresis, the region between BPB and XC was excised and washed in 1X Tris-borate pH 8.0 and EDTA (TBE) buffer. Gels were transferred to a wet nitrocellulose membrane in 1X TBE using Trans-Blot TurboTM Transfer System (1704150, Bio-rad) at constant 25 V for 20 min. The electroblotted membranes were briefly air-dried before crosslinking the RNA on the membrane using an ‘optimal crosslink’ in a UV crossliker (FB-UVXL-1000, Fisher Scientific). The membranes were pre-incubated in a hybridization buffer (0.9 M NaCl, 90 mM Tris-HCl, pH 7.5, 6 mM EDTA, 0.3% SDS, and 1% dry milk) at 37 °C for 1 h. Two DNA oligonucleotide probes, one targeting positions 18 to 36 of *E. coli* tRNA^Pro^(UGG) (5’-CCAAACCAGTTGCGCTACCA -3’) and the second targeting positions 19 to 36 of *E. coli* tRNA^Arg^(CCG) (5’-CGGAGGGCAGCGCTCTAT -3’) were each ^32^P-labeled at the 5’-end. These probes (each at 10^6^ cpm) were incubated with the pre-washed membrane in the hybridization buffer for 12 h while shaking. The probes were washed off by a 2xSSC buffer (0.3 M NaCl, 30 mM Na-citrate) at 37 °C two times, each time with mild shaking for 10 min. The membranes were then dried, and exposed to an imaging plate overnight. Imaging analysis was performed with a phosphorimager (Typhoon FLA 9500, GE) and the bands were quantified using ImageJ software (NIH). The charged fraction was calculated as the area encompassing the charged band divided by the sum of the charged and uncharged bands.

### qPCR analysis

WT and CRISPR-edited mutant cells were extracted for total RNA using TRIzol (Invitrogen) and Direct-zol RNA MiniPrep Kits (Zymo Research) according to the manufacturer instructions. It was processed to remove genomic DNA using RQ1 RNase-free DNase (M6101, Promega), followed by cDNA synthesis using RevertAid First Strand cDNA synthesis Kit (K1622, Thermo Scientific) according to the the manufacturer instructions. qPCR was carried out using the LightCycler® 96 Instrument. For each targeted gene, 18 µL of qPCR master mix containing 10 µL of SYBR Green I Master (04707516001, Roche), 7.6 µL of nuclease-free PCR grade water, 0.2 pmoles of gene-specific forward primer and 0.2 pmoles of gene-specific reverse primer was combined with 2 µL of cDNA template into each well of a 96-well plate (LightCycler® 480 Multiwell Plate 96, white, 04729692001, Roche). qPCR amplification was performed using the following cycling conditions: preincubation for 10 min at 95 °C, followed by 45 cycles of 3 steps of amplification with 10 sec at 95 °C, 10 sec at 48 °C, and 10 sec at 72 °C. The transcript level for each gene was determined using the standard curve generated from the control template by plotting the Cq values (y-axis) against the log initial concentration (x-axis).

The set of primers used for amplification of *ilvL* and *ilvG* genes are listed below: *ilvL* forward primer: 5′-ATGACAGCCCTT CTACG -3′; *ilvL* reverse primer: 5′-CTAAGCCTTTCCTCGTC-3′, *ilvG* forward primer: 5′-ATGAATGGCGCACAGTGG -3′; *ilvG* reverse primer: 5′-CCTGCTCATGTCGGCAT -3′. Normalization with idnT mRNA as a control confirmed the results.

### Reporter assay

The *leuL*-*leuA* genes from the *leu* operon was cloned from the *E. coli* MG1655 genomic DNA into pKK223-3 by NEBuilder HiFi DNA Assembly Cloning Kit (NEB) and the first 300 bp portion of *leuA* was translationally fused with the nano-Luc (nLuc) reporter gene. The four consecutive Leu codons CUA in *leuL* were mutated into the synonymous UUA codons by inverse PCR and self-ligation. The WT (CUA) and mutant (UUA) plasmids were transformed into *E. coli* JM109 *trmD-KO* cells maintained by human *trm5* (Masuda *et al*., 2019) together with pZS2R-mCherry plasmid for constitutive expression of mCherry for normalization. Cells were grown overnight in MOPS medium in the m^1^G37+ condition and inoculated 1:100 to fresh MOPS medium containing 0.1 mM IPTG and grown in m^1^G37+ and m^1^G37− conditions for 6 h at 37 °C. The nLuc readout was measured using Nano-Glo Luciferase Assay System (Promega), followed by normalization by the mCherry fluorescence level.

## Acknowledgements

The authors thank Chris Woolstenhulme and Ryuma Matsubara for preliminary experiments, Sean Moore and Ana Carr for materials to generate the *trmD-deg* strain, and Glenn Bjork for anti-TrmD antibodies. This study was funded by a JSPS fellowship to IM, NIH grants GM134931 and GM134931 to YMH, and GM110113 to ARB.

## Additional information

### Funding

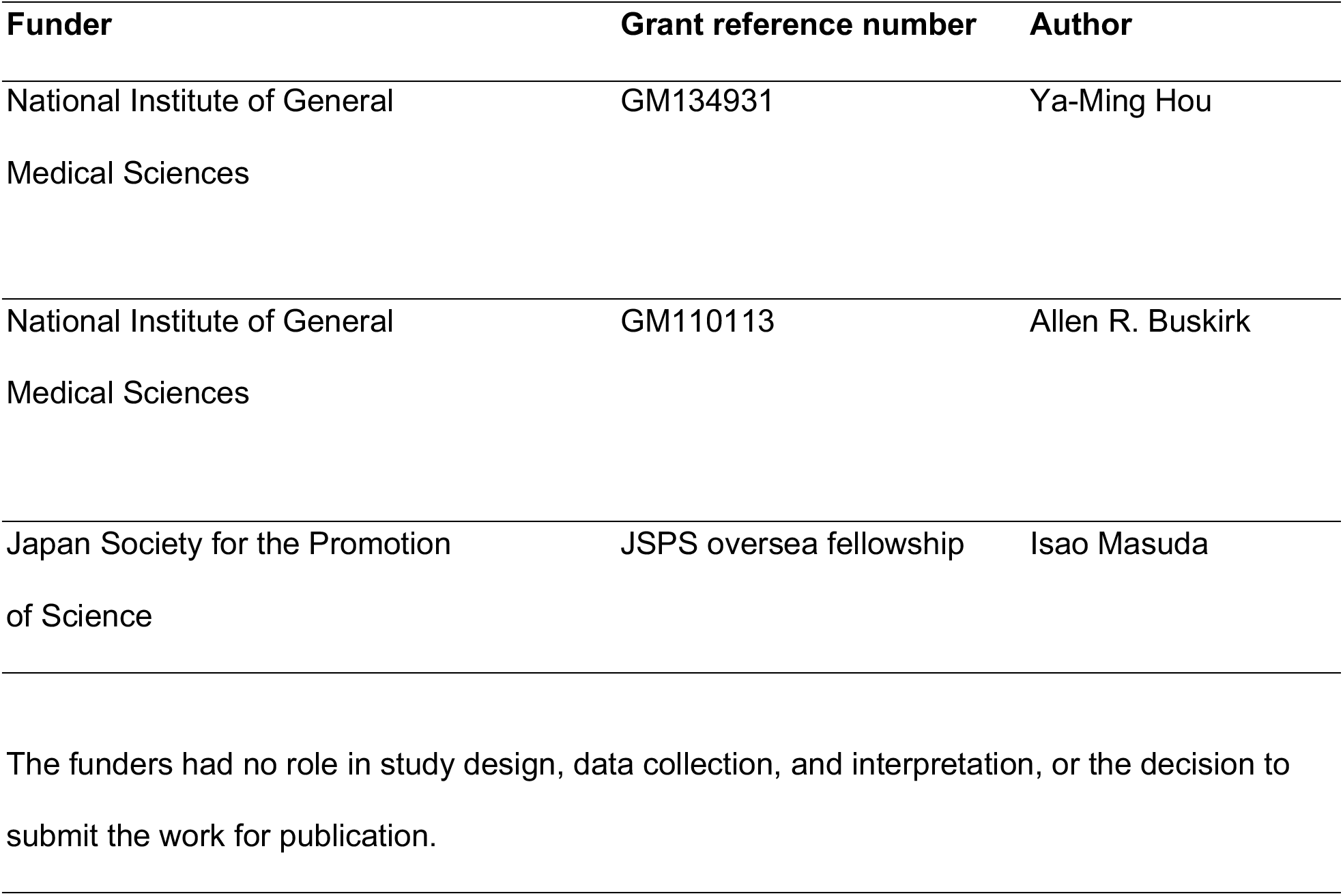

### Author contributions

Isao Masuda, Conceptualization, Formal analysis, Investigation, Methodology, Writing – original draft; Jae-Yeon Hwang, Conceptualization, Formal analysis, Investigation, Methodology; Tom Christian, Conceptualization, Formal analysis, Investigation, Methodology; Sunita Maharjan, Formal analysis, Investigation, Methodology; Fuad Mohammad, Methodology; Howard Gamper, Investigation, Methodology; Allen R. Buskirk, Conceptualization, Formal analysis, Supervision, Funding acquisition, Writing – review and editing; Ya-Ming Hou, Conceptualization, Formal analysis, Supervision, Funding acquisition, Writing – review and editing.

## Additional files

### Supplementary files

- Transparent reporting form

### Data availability

Sequencing data have been deposited in raw FASTQ files at the SRA and processed WIG files at the GEO under accession code GSE165592. Custom Python scripts used to analyze the ribosome profiling and RNA-seq data is freely available at https://github.com/greenlabjhmi/2021_TrmD

## Supplements

**Figure 1 Supplement.**
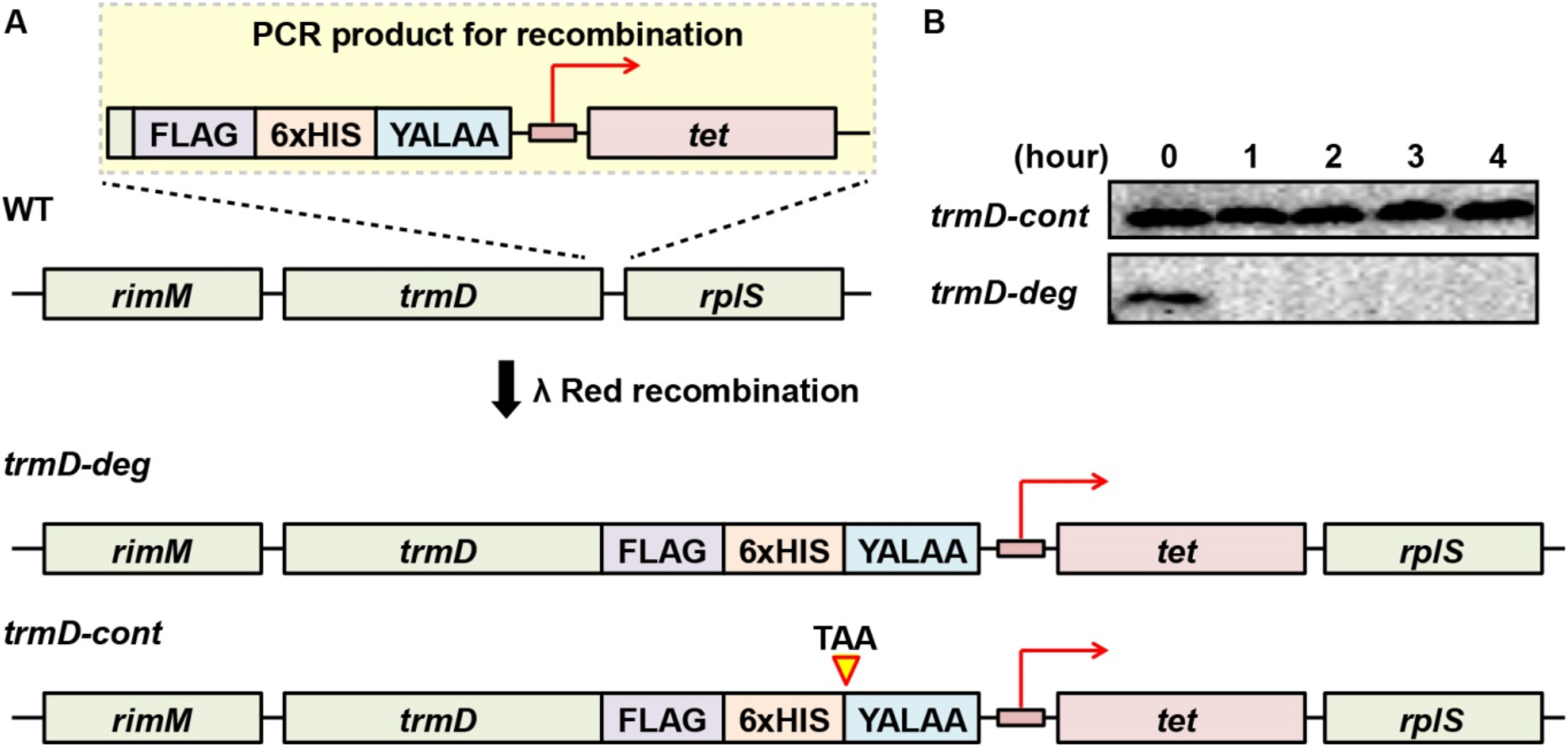
(A) Genomic contexts of *E. coli trmD-deg* and *trmD-cont* strains. A PCR product was amplified to contain sequences for a FLAG tag, a 6xHis tag, the degron peptide YALAA, and a tetracycline resistance marker with its promoter. This PCR product contained terminal extensions that were homologous to the 3’-end and the 3’-flanking region of *E. coli* chromosomal *trmD* and it was integrated into the chromosome *trmD* locus by λ Red recombination, generating the *trmD-deg* strain. To generate the *trmD-cont* strain, the same PCR product was used except that a stop codon TAA was added before the sequence for the YALAA tag. (B) Western blot analysis of TrmD protein over an extended time course. *E. coli trmD-deg* and *trmD-cont* strains were inoculated to LB + Ara (0.2%) grown at 37 °C for 4 h. Each culture was diluted to OD = 0.1 at the indicated time to maintain cells in the log phase and TrmD was probed by Western blot as in Figure 1D.

**Figure 3 Supplement.**
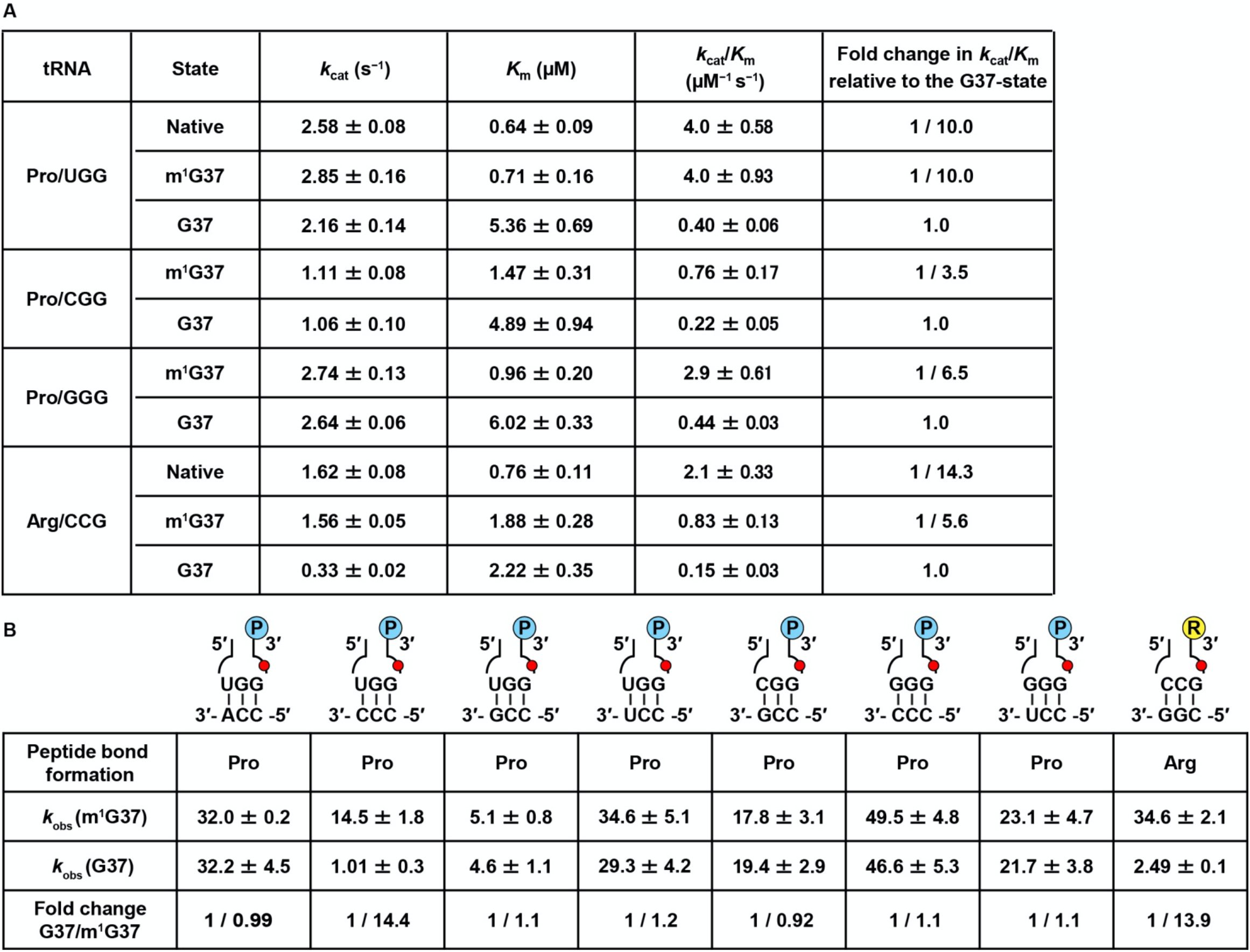
(A) Steady-state kinetic parameters *k*_cat_ (s^−1^) and *K*_m_ (µM) for aminoacylation of tRNAs. Each value is shown for the UGG, CGG, and GGG isoacceptors of *E. coli* tRNA^Pro^, and for the CCG isoacceptor of *E. coli* tRNA^Arg^, in the native-state, the m^1^G37-state, and the G37-state. The fold-change of *k*_cat_/*K*_m_ of the value in the native- or m^1^G37-state relative to the G37-state is shown for each isoacceptor. Errors are SD of three independent experiments (n = 3), and the data are presented as mean values ± SD. (B) Single-turnover *k*_obs_ values for peptide-bond formation of tRNAs. Each value is shown for the UGG, the CGG, and the GGG isoacceptors of *E. coli* tRNA^Pro^, and for the CCG isoacceptor of *E. coli* tRNA^Arg^, in the m^1^G37-state and the G37-state. The codon-anticodon pair tested is indicated in the cartoon on the top of each column. The fold-change of *k*_obs_ of the value in the m^1^G37-state relative to the G37-state is shown at the bottom. Errors are SD of three independent experiments (n = 3), and the data are presented as mean values ± SD.

